# Selective covalent-allosteric tools to dissect Akt2

**DOI:** 10.64898/2026.06.15.732341

**Authors:** Lena Quambusch, Danilo D’Angelo, Tonia Kirschner, Maria Beerbaum, Laura Depta, Fabian Schnecke, Janina Niggenaber, Sven Brandherm, Jörn Weisner, Matthias P. Müller, Leif Dehmelt, Daniel Rauh

**Affiliations:** Faculty of Chemistry and Chemical Biology, TU Dortmund University and Drug Discovery Hub Dortmund (DDHD), Zentrum für Integrierte Wirkstoffforschung (ZIW), Otto-Hahn-Strasse 4a, 44227 Dortmund, Germany; L.Q.: Department of Biochemistry, University of Cambridge, 80 Tennis Court Road, CB2 1GA, Cambridge, UK

## Abstract

The protein kinase Akt and its isoforms play a crucial role in various diseases. Unique functions of the individual isoforms (Akt1, Akt2, Akt3) might be essential for survival in malignancies. Particularly for Akt2, it was reported that a knock-out led to diabetic phenotype and might be correlated with clinically adverse hyperglycemic effects observed in pan Akt-treatment. Enduring failure of Akt inhibitors in the clinic indicates the necessity for a thorough understanding of the underlying biology, preferably by using highly isoform-selective small molecules. Here we report the structure-guided development of Akt2-selective covalent-allosteric probe molecules, that can be successfully modified within a complex environment using biorthogonal chemistry. Thus, enabling first Akt2-specific pull-down studies and the use in functional studies, such as selective fluorescent labeling in cellular systems. These chemical probes expand our toolbox to dissect the critical questions of Akt2’s function in health and disease, thereby paving the way for novel therapeutic strategies based on thorough mechanistic insights.

## Introduction

As a central mediator within the PI3K/Akt/mTOR-pathway, the protein kinase Akt can be linked to various diseases such as cancer and diabetes.^1–3^ The human genome encodes three isoforms (Akt1, Akt2, Akt3), that share a high overall sequence homology. Their differing functions are presumably determined through intracellular localization and tissue-specific expression levels.^4^ Whereas Akt1 is ubiquitously expressed, Akt2 is enriched in muscle tissue and specifically within mitochondria, while Akt3 displays higher expression levels in neurons.^5^ First functional studies in knockout mice revealed insights into the individual isoform functions. Those invasive experiments showed a link between Akt1 to proliferation and antiapoptotic behavior.^6^ In contrast, Akt2 deletions illustrate a type-2 diabetic phenotype along with impaired glucose uptake.^7,8^ Finally, neuronal malfunctions can be connected to the absence of Akt3 and alterations within the fatty acid metabolism.^9,10^

More recently, genetically modified cancer models were created that put previous results for the roles of individual isoforms into question, indicating that invasive experiments with the aim to dissect the specific contributions of the Akt isoforms towards this proliferative disease have limitations.^11,12^ The aforementioned knockout studies in aggressive breast cancer models imply that Akt2 is involved in metastasis, whereas in lung cancer models, Akt1 indicates tumor-initiating behavior. In both scenarios, a different isoform-selective therapeutic strategy would be needed to impair their individual function.

For a more suitable elucidation of the function and biological role of Akt2, less invasive perturbations, preferably mediated by highly selective bioactive small molecules, are needed, which will facilitate concentration-dependent and temporally controllable experiments.^13^

Recently, we reported on Akt2 isoform-selective covalent-allosteric inhibitors that retained their profile within cellular model systems and qualified as potent molecules to perturb the isoform for functional studies.^14,15^ Those molecules represent an optimal basis for generating functionalized probes that facilitate an in-depth dissection of biological functions. The introduction of biorthogonal groups, such as alkynes, for subsequent click-chemistry and proteomic analysis, will enable a broad range of follow-up experiments.^16,17^

This study presents a structure-guided design of highly functionalized covalent-allosteric probe molecules to address Akt2 within complex systems. Our design emerged from an Akt1 hybrid X-ray structure containing allosteric site mutations mimicking the corresponding Akt2 pocket. Synthesis of alkyne probes followed by biochemical evaluation led to conserved selectivity profiles and allowed biorthogonal chemistry on covalently labeled Akt2 in cell lysates. Subsequent pull-down studies revealed the selective enrichment of Akt2 in complex environments and the probes usage in functional studies such as fluorescence microscopy.

## Results

Over the course of these studies, a few full-length Akt2 structures have been reported, including a publicly available structure (PDB: 9C1W),^18,19^ yet these became accessible only after the initiation of our work. For this reason, we have previously worked with homology models of Akt2 on the basis of Akt1 structures for our docking and design approaches.^14^ However, in order to further substantiate our previous findings, our first aim within this study was to obtain an Akt2-mimicry crystallization system based on the highly reliable and reproducible Akt1 crystallization construct. Encouraged by our previous results and with the obvious differences found via sequence alignments, we introduced several amino acid mutations within the allosteric pocket of Akt1 via site directed mutagenesis (S205T, D262E, K268R, N269D and deletion of E267).^20,21^ Similar approaches were reported before, e.g., factor XA pocket into another protease.^22^ This resulted in an Akt2-mimicry construct and expression and purification of this novel hybrid protein worked sufficiently well and allowed various soaking experiments and X-ray screenings.

In complex with the reversible allosteric inhibitor miransertib, the newly designed mimicry construct resulted in well-diffracting crystals and a complex structure (see Fig. 1B).^23^ The structure reveals common protein-ligand interactions of this type of inhibitor. In direct comparison to an Akt1 complex, it shows an altered αE-helical structure around Arg267 due to the deletion of one residue. This confers with former homology models and reported Akt2 kinase domain complexes.^14^ The quite flexible side chain of the basic amino acid Arg267 is not resolved in this structure. Compared to several published Akt1 structures, the corresponding lysine residue at this position is always well resolved. Once again, this indicates an increased flexibility of the amino acid in this hybrid construct and emphasizes an altered size of the allosteric Akt2 binding pocket at this solvent-accessible site.

**Fig. 1:**
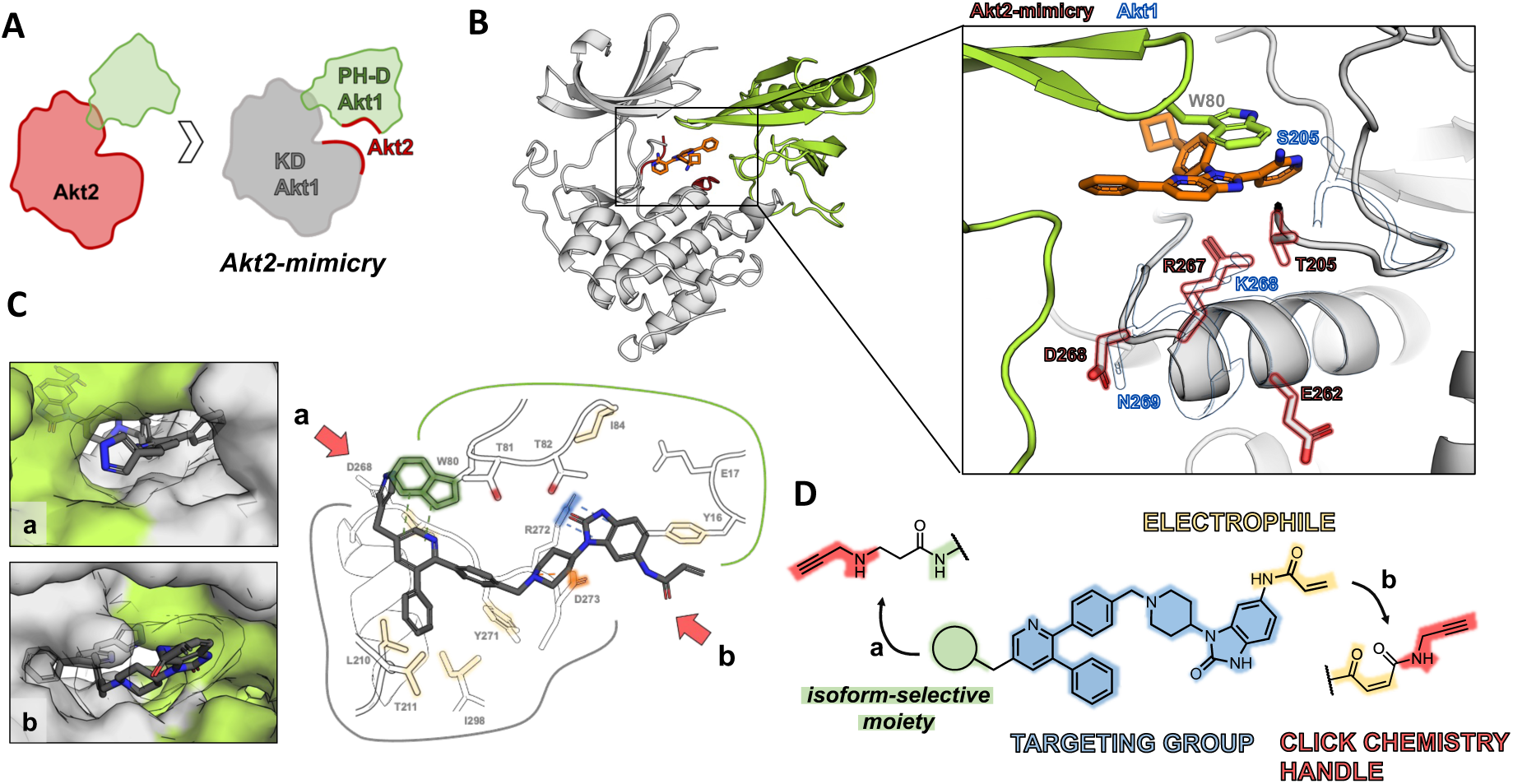
Design of Akt2-selective covalent-allosteric probe molecules based on structural insights from an allosteric Akt2-pocket mimicry construct. A: Schematic representation of the developed Akt2-mimicry construct for protein crystallography, B: Complex structure of Akt2-mimicry construct with allosteric inhibitor miransertib (PDB: 29mj). Close-up shows mutated amino acid side chains (red) and the corresponding *α*E-helix in Akt1 (blue) PDB: 6s9w;^15^ C: Docking studies of isoform-selective CAAI 14 into Akt2-mimicry construct highlights interactions (green: *π*-*π*-Stacking, yellow: hydrophobic interactions, blue: cation-*π*-interaction, orange: salt bridge). The surface representation on the right shows the two-sided solvent accessible allosteric pocket (a,b), D: Schematic design of covalent probe molecules either modified with a clickable alkyne moiety at the isoform-selective residue of the ligands (a) or at the electrophilic warhead (b).

On the basis of the new complex crystal structure, a docking study with the recently reported Akt2-selective CAAI **14** was performed (see Fig. 1C).^14^ Again, the docking pose highlights various interactions between the CAAI and amino acid side chains in the allosteric pocket. Mainly this interaction is mediated through *π*-*π*-stackings, hydrophobic interactions, and an important salt bridge between the tertiary amine of the piperidine ring system and Glu273.

Furthermore, the *in silico* study shows two different solvent accessible sides of the addressed allosteric pocket - one at the site of the selective moiety in the western part of the inhibitor and another at the eastern part of the molecule close to the electrophilic warhead - indicating possible exit vectors to introduce an alkyne moiety as a biorthogonal handle. In former studies, several molecules were identified that possess a distinct selectivity profile towards Akt2 while sparing the other two isoforms, Akt1 and Akt3 (SI-Fig. 1).^14,15^ Whereas the pyrazinone-based inhibitor **16b** and the pyridine **14** are not suitable for a chemical modification of the selectivity-mediating moiety in the western part of the molecule, those ligands are optimal for the introduction of an altered warhead to allow the connection of an alkyne. In contrast, pyridine **17** with *meta*-aniline as a selectivity group can be used to introduce an alkyne at this site of the allosteric pocket, which would have the advantage of not altering the warhead.

For the transformation of the described CAAIs into molecular probes, their synthetic routes were slightly altered. The introduction of a malonic acid group as an electrophilic warhead while giving the opportunity to modify the terminal acid moiety should be included in the last step of the synthesis for the **14** and **16b**-based probes (Scheme 1B). Within the reported synthesis of **16b** the last step was the introduction of the acrylamide, hence the prior intermediate **14b** could be used immediately for further modification.^15^ To get the anilinic version of CAAI **14,** the benzimidazolone part of the molecules needed to be Boc-protected before introducing the boronic ester. The reductive amination led to molecule **1**. Followed by Suzuki coupling with reported pyrazole pyridine **S22** and deprotection of the aniline yielded molecule **4**. From here, both reversible precursors **4** and **14b** were coupled with maleic anhydride to give the corresponding acids **5a** and **5b**. As the last step, the acids were coupled with propargyl amine to give the finals **probe 1** and **probe 2**.

**Scheme 1.**
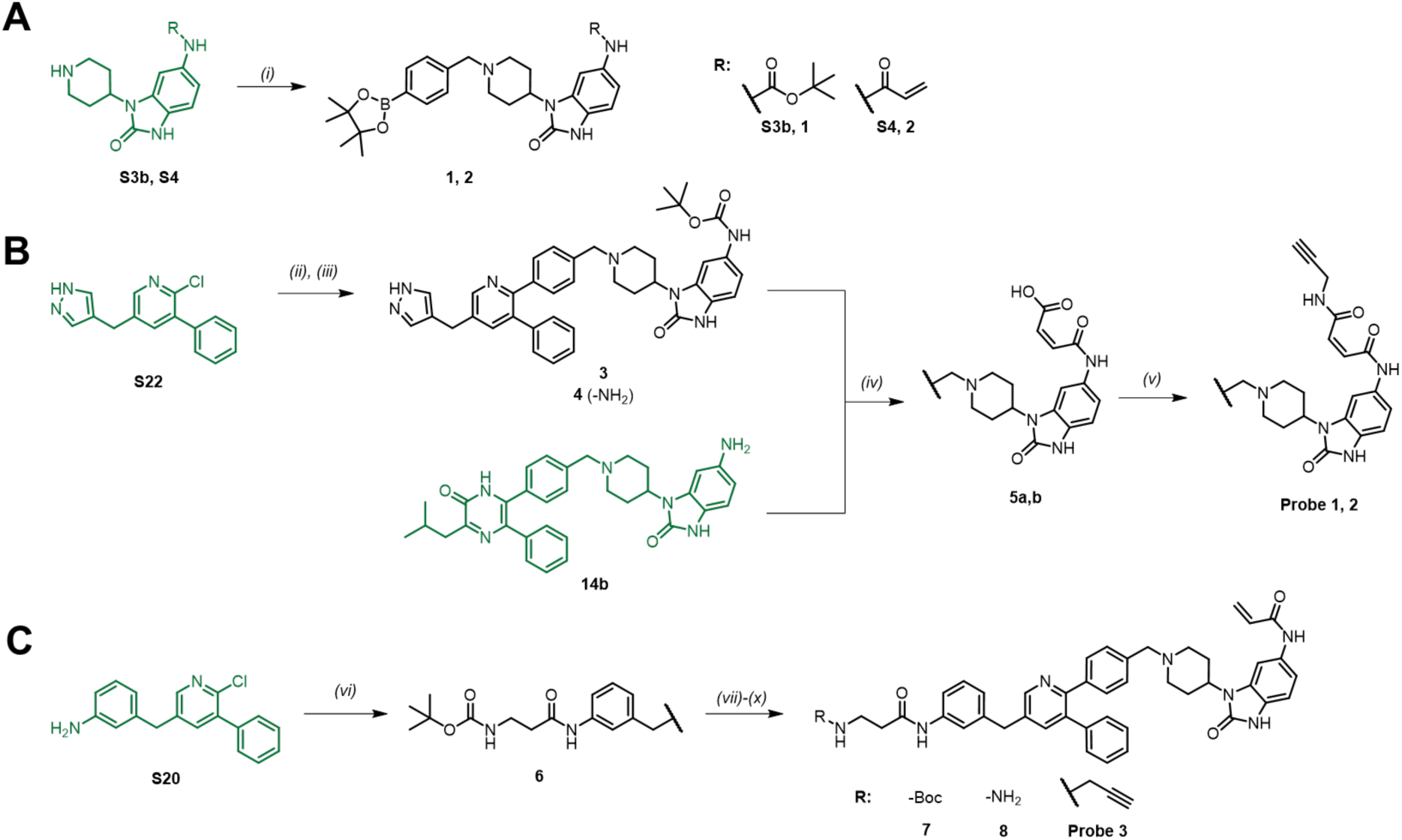
**Synthesis scheme Akt2-selective covalent-allosteric probe molecules.*^a^*** *^a^*Reagents and conditions: (*i*) 4-Formyl phenylboronic pinacol ester, NaCNBH_3_, Et_3_N/AcOH, MeOH, 75 °C, 12 h (30-66 %); (*ii*) Pd(PPh_3_)_2_Cl_2_, K_2_CO_3_, 1,4-dioxane/H_2_O (5:1), 130 °C, *µ*w, 2 h (48 %); (*iii*) DCM/TFA (3:1), rt, 16 h; (*iv*) Maleic anhydride, DCM, 16 h, (89 -91%); (*v*) Propargylamine, DCC, THF, 6 h, (51-58 %); (*vi*) HATU, DIPEA, MeCN, rt, 16 h, (73 %); (*vii*) Pd(PPh_3_)_2_Cl_2_, K_2_CO_3_, 1,4-dioxane/H_2_O (5:1), 130 °C, *µ*w, 2 h (29 %); (*viii*) DCM/TFA (3:1), rt, 12 h (98 %); (*ix*) Propargyl bromide, K_2_CO_3_, DMF, rt, 3 h, (30 %).

The modification of CAAI **17** to result in **probe 3** needed the introduction of a short linker to ensure that the alkyne would be solvent-accessible. Therefore, *meta*-aniline **S20** was coupled to β-Boc-alanine to give molecule **6**.^14^ This was coupled to the boronic ester **2** via Suzuki cross-coupling (**7**) and then deprotected with TFA to give intermediate **8**. At the end, the primary amine was alkylated with propyne bromide in a nucleophilic substitution and gave **probe 3**. Three novel alkyne probes were generated based on well-characterized CAAIs (see Fig. 2A).

**Fig. 2:**
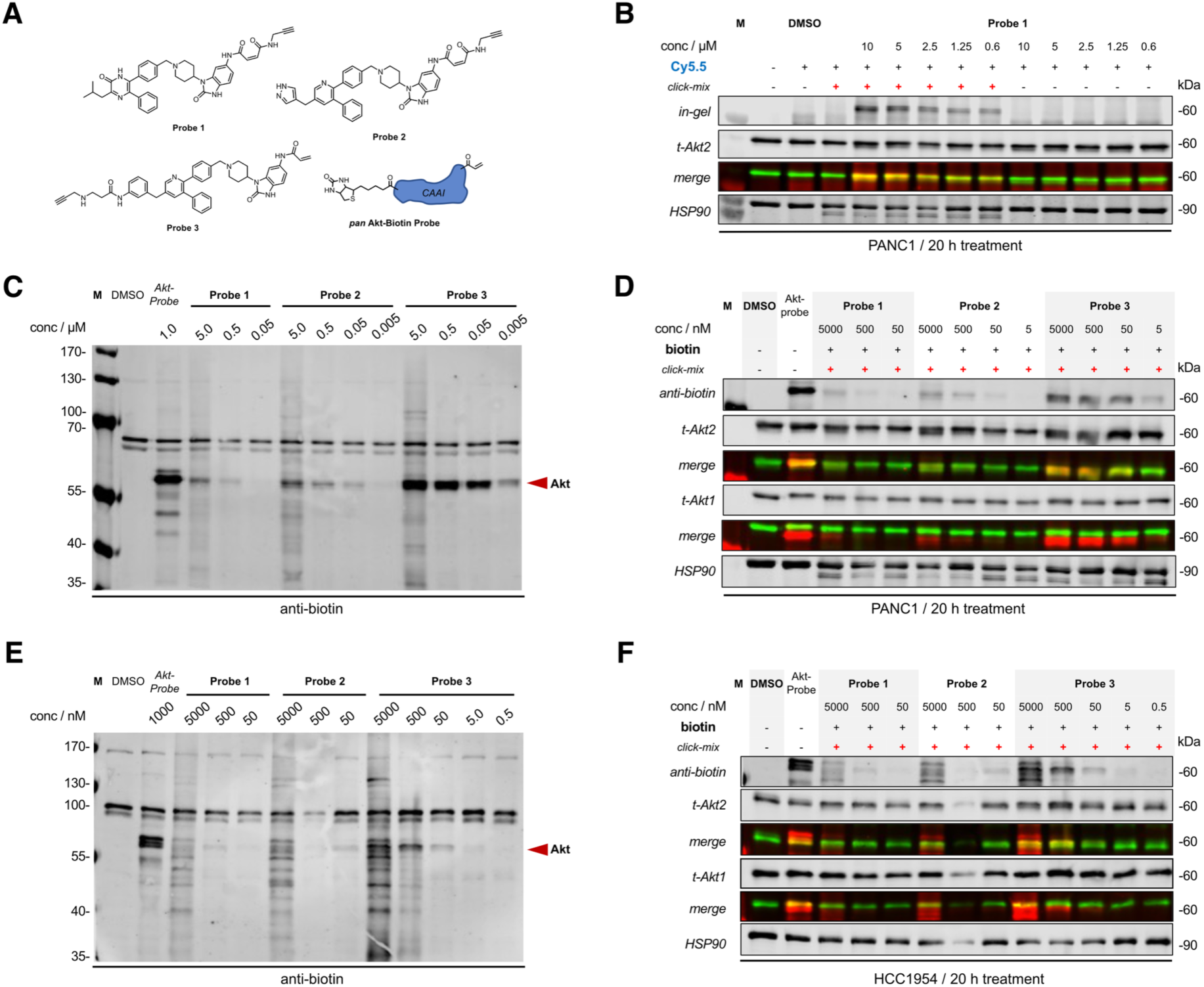
Covalent-allosteric alkyne probes selectively modify Akt2 in treated cancer cells. A: Structures of newly synthesized molecular alkyne probes (probe 1-3) and a schematic representation of a CAAI-based *pan*-Akt molecular probe pre-labeled with biotin already; B: Immunoblots showing Cy5.5 fluorescence and tAkt2-signal after 20 h treatment of PANC1 cells with **probe 1**. Different concentrations ranging from 10 to 0.6 µM were used and the resulting lysates exposed to Cy5.5-azide and copper-containing or copper-free click-mix; C: Overview of an anti-biotin-stained immunoblot of PANC1 pancreatic cancer cells treated with different concentrations of all CAAI probes followed by click-chemistry modification with biotin-PEG3-azide. D: Compiled immunoblot of the experiment shown in C but stained with both tAkt1- and tAkt2 antibodies to access the isoform-selective labeling of the probe-treated PANC1 cells. E: Overview of an anti-biotin-stained immunoblot of HCC1954 breast cancer cells treated with different concentrations of all CAAI probes followed by click-chemistry modification with bio-tin-PEG3-azide; F: Compiled immunoblot of the experiment shown in E but stained with both tAkt1- and tAkt2 antibody to access the isoform-selective labeling of the probe treated HCC1954 cells.

In order to assess the impact of further groups introduced into the molecules with respect to the selectivity profile, the small set was evaluated *via* HTRF kinEASE activity assay (Tab. 1). The data shows that it was possible to retain the overall selectivity ratios towards Akt2 for every probe. Modifications of the warhead through the maleic electrophile moiety entail the potency of the probes compared to the parental versions. This could be due to a different orientation of the elongated warhead in the pocket, a restricted flexibility to the two addressed cysteine residues, or the electrophilic nature of this *Z*-configurated Michael-acceptor system might slow the overall covalent bond formation.^24^

**Table 1:**
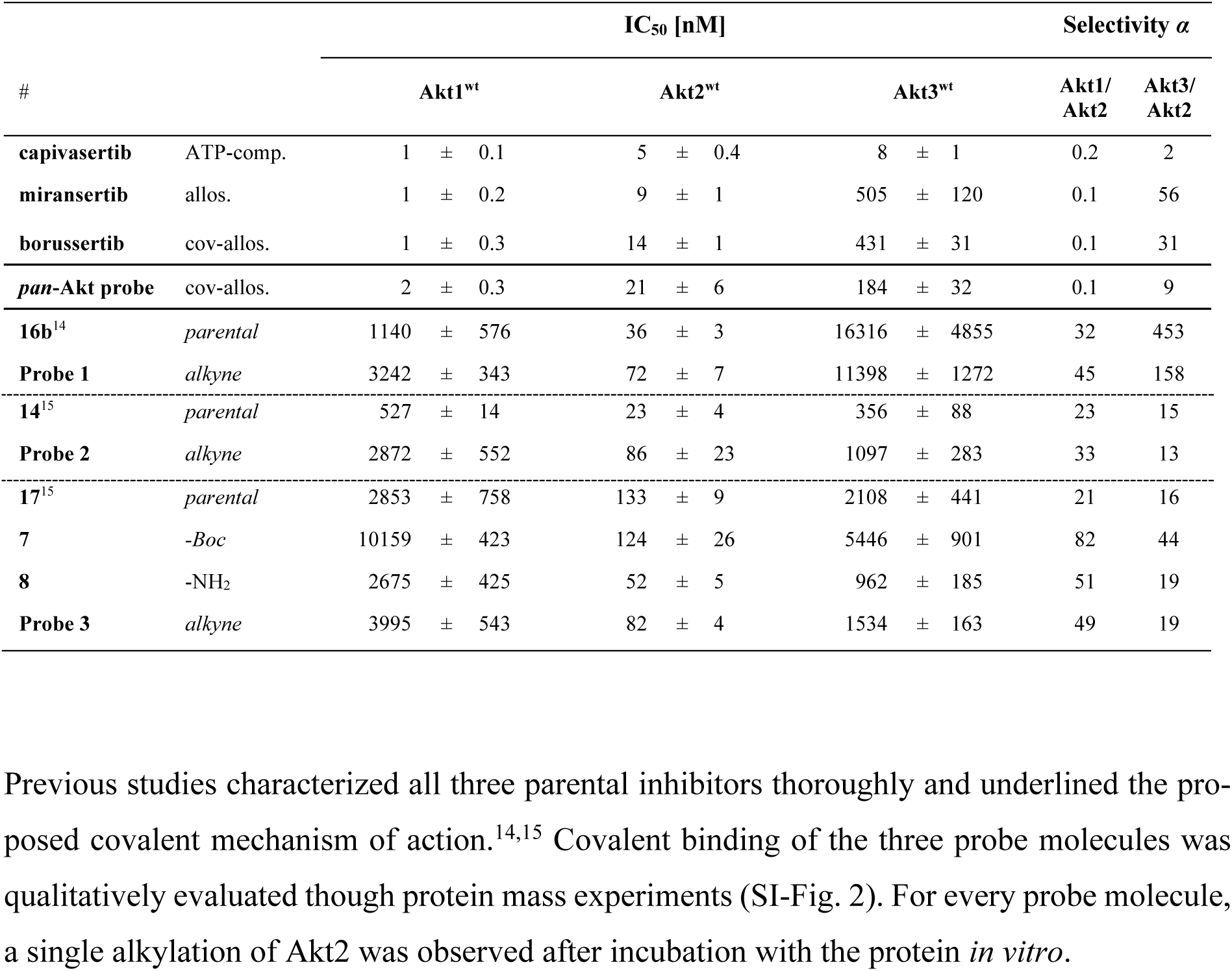
Biochemical evaluation of novel covalent-allosteric Akt probes with the three Akt isoforms.

| # | | IC<sub>50</sub> [nM] | | | Selectivity $\alpha$ | |
| --- | --- | --- | --- | --- | --- | --- |
|  |  | Akt1<sup>wt</sup> | Akt2<sup>wt</sup> | Akt3<sup>wt</sup> | Akt1/<br>Akt2 | Akt3/<br>Akt2 |
| **capivasertib** | ATP-comp. | 1 ± 0.1 | 5 ± 0.4 | 8 ± 1 | 0.2 | 2 |
| **miransertib** | allos. | 1 ± 0.2 | 9 ± 1 | 505 ± 120 | 0.1 | 56 |
| **borussertib** | cov-allos. | 1 ± 0.3 | 14 ± 1 | 431 ± 31 | 0.1 | 31 |
| **pan-Akt probe** | cov-allos. | 2 ± 0.3 | 21 ± 6 | 184 ± 32 | 0.1 | 9 |
| **16b**<sup>14</sup> | *parental* | 1140 ± 576 | 36 ± 3 | 16316 ± 4855 | 32 | 453 |
| **Probe 1** | *alkyne* | 3242 ± 343 | 72 ± 7 | 11398 ± 1272 | 45 | 158 |
| **14**<sup>15</sup> | *parental* | 527 ± 14 | 23 ± 4 | 356 ± 88 | 23 | 15 |
| **Probe 2** | *alkyne* | 2872 ± 552 | 86 ± 23 | 1097 ± 283 | 33 | 13 |
| **17**<sup>15</sup> | *parental* | 2853 ± 758 | 133 ± 9 | 2108 ± 441 | 21 | 16 |
| **7** | -Boc | 10159 ± 423 | 124 ± 26 | 5446 ± 901 | 82 | 44 |
| **8** | -NH<sub>2</sub> | 2675 ± 425 | 52 ± 5 | 962 ± 185 | 51 | 19 |
| **Probe 3** | *alkyne* | 3995 ± 543 | 82 ± 4 | 1534 ± 163 | 49 | 19 |

In the case of **probe 3** as well as for the intermediates of the synthesis, an improvement in potency was observed, which might originate from additional contacts to the protein. These results indicate that this approach might be generally superior since the warhead and the covalent reaction are unaltered and thus cannot negatively affect the potency.

Previous studies characterized all three parental inhibitors thoroughly and underlined the proposed covalent mechanism of action.^14,15^ Covalent binding of the three probe molecules was qualitatively evaluated though protein mass experiments (SI-Fig. 2). For every probe molecule, a single alkylation of Akt2 was observed after incubation with the protein *in vitro*.

In order to assess if the generated alkyne probes can be used as tools for click-chemistry, cellular treatment of PANC1 cells was conducted (Fig 2B). The relatively moderate concentrations (> 600 nM) of **probe 1** were used in this setup. Generated lysates were then used for click-reaction with Cy5.5-azide to result in a fluorescence signal. A clear concentration-dependent signal was observed around the size of Akt within samples containing copper, azide, and alkyne-treated cells. DMSO control samples and copper-free lysates did not show any fluorescence signal. A co-stain with tAkt2 underlines the selective labeling of the addressed target in PANC1 cells nicely through an overlay of reporter and antibody signal, whereas Akt1 was not labeled, again highlighting the very good overall selectivity.

Especially for further functional studies and enrichment of Akt2, the modification with biotin is of important interest and was evaluated after 20 h treatment of PANC1 cells with all three probes and an already biotin-tagged *pan*-Akt CAAI as control (Fig. 2C). The SA-DyLight™ stained blot highlights present endogenously biotinylated proteins within the PANC1 cells and gives constant signals at the various concentrations of the probes.^25^ The biotin-tagged pan-Akt probe visualizes the overexpressed fraction of Akt2 in those cells and some other small modifications. Further, it seems **probe 2** can label Akt even at concentrations as low as 50 nM, whereas **probe 1**’s limit seems to be around 500 nM, as seen in Fig. 2C. In contrast, **probe 3** treated cells correspond to a similar portion of modified Akt as the pan-Akt biotin-probe. There is a clear and strong signal visible, and concentrations between 500 and 50 nM give no off-target labeling, whereas at 5 µM probe concentration, faint unspecific signals appear.

The selective modification of Akt2 in PANC1 cells was also validated by staining for tAkt1 (Fig. 2D). Based on the pan-Akt biotin probe, it’s clear that only a very small portion of Akt1 is expressed in PANC1, which is located above the prominent Akt2 band and can be co-stained. None of the herein-generated alkyne probes seem to significantly label Akt1 and they only result in a single biotin signal corresponding to Akt2.

Since the PANC1 cell line is somewhat biased to a selective Akt2 modification due to Akt2 overexpression, further assessment of the selective properties was also assessed in the HCC1954 cell line that does not overexpress any of the three Akt isoforms (Fig 2E).^26^ The pan-Akt biotin probe results in three clear signals that presumably correspond to the Akt Isoforms and confirms the equivalent expression of those kinases within this particular breast cancer cell line, which is in good agreement with the isoform specific signal of Akt1 and Akt2 (Fig 2F). Alkyne **probe 1** and **probe 2** show a few modifications across the lysate, but the signal for Akt2 is not dominantly present, even at high concentrations. In the case of **probe 3**, the biotin signal nicely corresponds to the Akt2-labeled protein (Fig. 2E). Concentrations around 500 nM and below give very limited off-target modifications. Hence, resolving the isoform-selective labeling in this cell line is possible, but the overall efficacy is impaired in contrast to the PANC1 cell line.

Whether the alkyne probes can selectively modify Akt2 within a complex cell lysate, concentration-dependent treatment of PANC1 cells was investigated (Fig. 3A/B). The results underline a higher off-target modification at 5 µM compared to the treated cells (Fig. 2). Surprisingly even lower concentrations of **probe 2** can selectively modify Akt2 in this experimental setup, whereas this molecular probe was limited in labeling ability used in treatments before. In all three scenarios, a concentration around 500 nM or lower seems to be an effective amount to detect distinct labeling of Akt2 while sparing the other two Akt isoforms. **Probe 3**, on the other hand, is weaker in modification if concentrations are below 50 nM, which might correlate to the actual IC_50_-value (80 nM, Tab. 1).

**Fig. 3:**
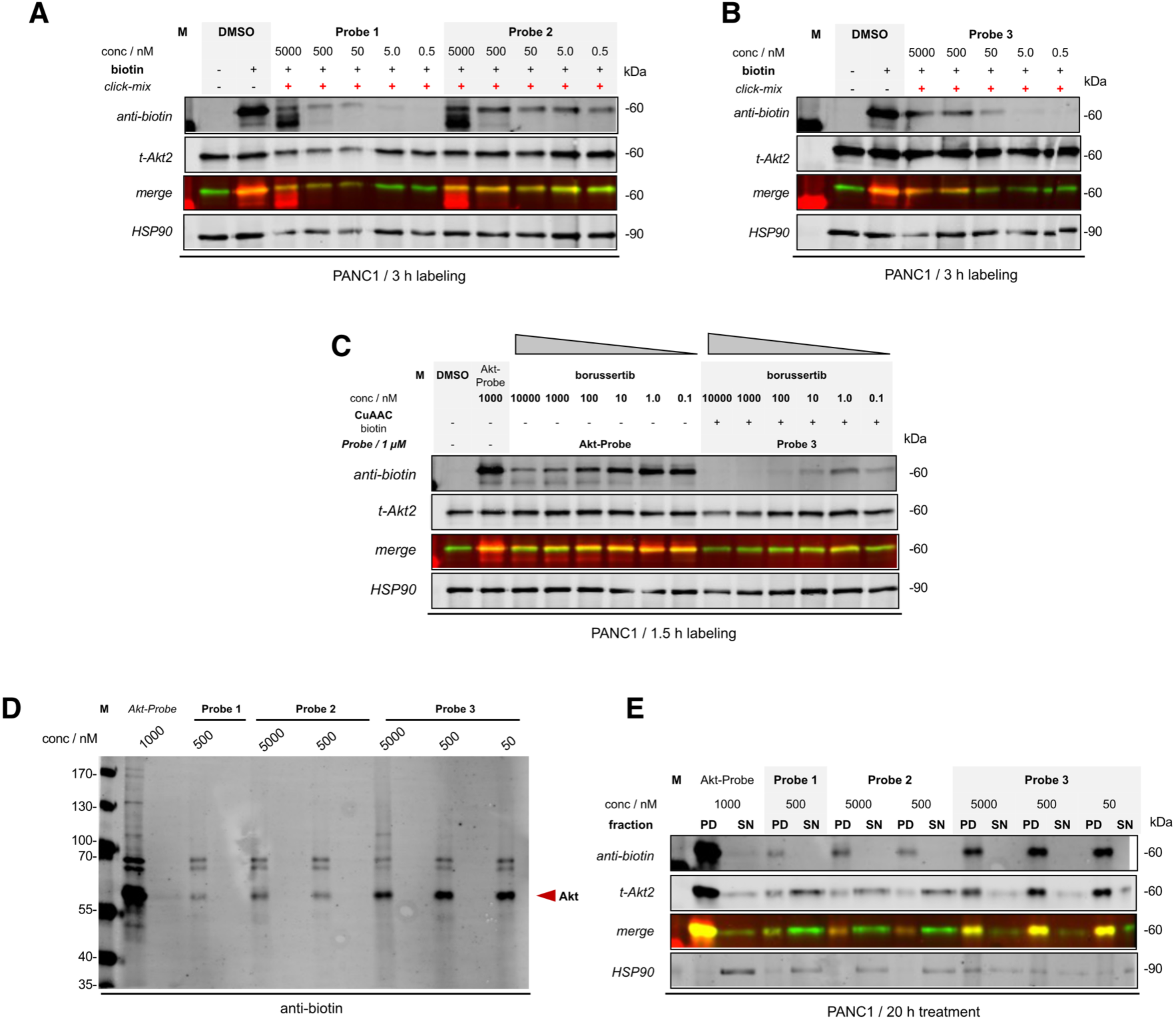
Covalent-allosteric alkyne probes selectively modify and enrich Akt2 in a complex environment. A/B: Overview of compiled immunoblots of PANC1 cell lysate treated with different concentrations of **probe 1**, **probe 2** and **probe 3** followed by click-chemistry modification with biotin-PEG3-azide. Blots were stained with both tAkt1- and tAkt2 antibodies to assess the isoform-selective labeling; C: Immunoblot of PANC1 lysates treated with different concentrations of borussertib for 3 hours followed by the addition of 1 µM of the *pan*-Akt probe or **probe 3**. The biotin signal increases with lower borussertib concentrations as the allosteric pocket is preoccupied; D: Overview of a biotin-stained immunoblot of PANC1 biotin-streptavidin pull-down experiments treated with different concentrations of all probes after modification *via* click-chemistry (PD: pull-down, SN: supernatant); E: Compiled immunoblot of the experiment shown in E but stained with both tAkt1- and tAkt2 antibodies to assess the isoform-selective labeling of the pull-down and supernatant fractions.

Further, a competition experiment with covalent-allosteric Akt inhibitor borussertib was performed to assess whether Akt can be preoccupied with another CAAI and therefore is impaired to be labeled with a molecular probe (Fig. 3C).^27^ As a comparison, the pan-Akt biotin probe was used in the same setting. In the case of **probe 3**, the experiment highlights an increasing signal when the concentration of borussertib is lowered, and no modification can be observed at the highest inhibitor concentration. This experiment validates the occupation of the targeted binding pocket of Akt2, which is preoccupied if a potent CAAI such as borussertib was used before.

To enrich the target protein, Akt2 pull-down experiments with all three probes and treated PANC1 cells were performed (Fig. 3D). Again, the pan-Akt biotin probe was used as a positive control. The immunoblot shows enrichment of biotinylated proteins around 70 kDa and another single protein around 60 kDa. For **probe 1** and **probe 2** are less intense signals visible at 60 kDa than for **probe 3**. However, even with decreasing alkyne concentration, **probe 3** is able to accumulate a similar amount of protein within the pull-down fraction. Co-staining with an anti-Akt2 antibody validates the specific enrichment within the PD fraction and detects higher amounts of unmodified Akt2 in the supernatant when the concentration of **probe 3** decreases (Fig. 3E). In contrast, **probe 1** and **probe 2** are not able to enrich the same amounts of Akt2. The control biotin-probe enriches other proteins at the used concentration of 1 µM. Overall, underlining the previous experiments’ observations, **probe 3** is more suitable for effectively modifying Akt2 in complex systems, even at a concentration of around 50 nM.

As a final functional complementation and to prove the feasibility of covalent-allosteric inhibitors as selective tools, we used HEK293 cells and investigated the in-cell clickability of **probe 3**. To assess the selectivity of the clickable alkyne probe toward Akt2, HEK293 wild-type (WT) and HEK293 Akt2 knock-out (KO) cells were treated with the alkyne-containing Akt2 inhibitor **probe 3**. Following treatment, cells were fixed and subjected to a copper-catalyzed click reaction to conjugate a Cy3–azide fluorophore to the alkyne probe. Fluorescence analysis revealed a substantially higher Cy3 signal in WT cells compared to AKT2 KO cells, although a residual signal was detectable in the KO condition (see Fig. 4). This observation was further supported by pull-down studies using biotin-azide with probe 3 in both cell lines, which revealed nonspecific labeling of a smaller protein alongside only modest Akt2 enrichment (see SI-Fig. 3). This off-target labeling likely accounts for the residual fluorescence signal observed in KO cells. Transfection of an Akt2–GFP construct into Akt2 KO cells restored probe labeling, predominantly in cells that expressed Akt2-GFP. Together, these results demonstrate that cellular labeling by the alkyne probe is largely Akt2 level-dependent, supporting its selectivity in cells. Importantly, the clickable nature of the probe enables versatile downstream applications, including fluorescence-based visualization of Akt2 localization and engagement, providing a valuable tool for future studies of Akt2 distribution and function in different cellular contexts.

**Fig. 4:**
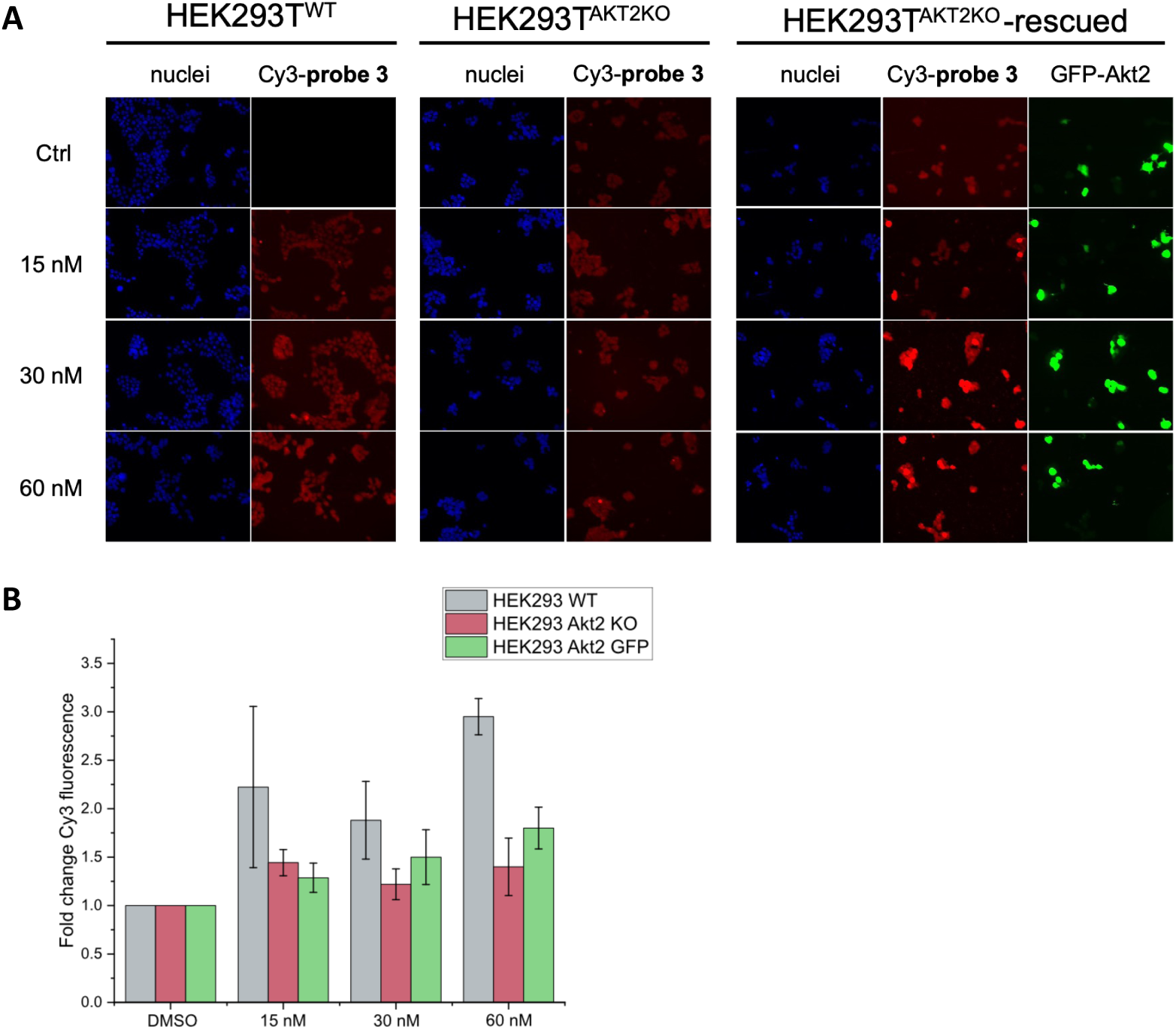
Akt2-selective probe can be fluorescently labeled in HEK293 cells for microscopy studies. A: Representative fluorescence microscopy images of parental HEK293T cells, HEK293T Akt2 knockout (Akt2 KO) cells, and Akt2 KO cells transiently rescued by re-expression of GFP-Akt2. Cells were left untreated (DMSO) or treated with 15 nM, 30 nM, or 60 nM of **probe 3**, followed by click chemistry conjugation with azide–Cy3 for fluorescent detection. Images are representative of at least 3 independent experiments; B: Quantification of intracellular Cy3-**probe 3** signal. For each independent experiment, the median fluorescence intensity per condition was calculated, and the graph represents the mean ± SD of these median values from 3 independent experiments.

## Discussion

Multiple parameters of the inhibitors themselves and their behavior in the complex environment play a crucial role in choosing the right tool for labeling experiments.^28^ In our study, we presented two different approaches introducing an alkyne moiety within established covalent-allosteric inhibitors. One is based on a modified warhead and a possibility of interfering with the alkylation step when the CAAIs bind to the targeted cysteines. For another design, where the warhead is not altered, the opposite solvent-exposed site of the allosteric pocket was utilized to introduce an alkyne moiety. The influence of linkers and distances of biorthogonal groups are widely accepted to have a tremendous impact on the functionality of the designed probes and chemical tools (e.g., degraders, bifunctional molecules).^29^

The experimental setup and question can direct the use of the here presented probe molecules. For example, whereas **probe 3** is characterized by a very strong enrichment and modification of Akt2 when used in treatment experiments, **probe 2** might be the tool of choice if lysed cells needed to be screened for Akt2. Upon assessing all three probes across different cell lines, **probe 3** emerged as the most efficient. The differential performance observed suggests that **probe 1** and **probe 2** may be less effective in certain cellular environments. Alternatively, the enhanced efficacy of these alkyne-labeled probes in PANC1 cells could be attributed to elevated Akt2 expression levels in this pancreatic cancer line, which may promote more stoichiometric binding and labeling.

To validate the utility of covalent-allosteric inhibitors as chemical tools for target identification, we evaluated the cellular applicability of **probe 3** in live cells and complex proteomes. In-cell click experiments in HEK293 cells demonstrated concentration-dependent fluorescent labeling that was significantly enhanced compared to DMSO controls. Interestingly, comparable labeling was observed in HEK293 Akt2 KO cells, suggesting off target labeling in this specific cell line. These findings confirm that probe 3 exhibits preferential labeling of Akt2, though the modest labeling efficiency and minor nonspecific interactions observed suggest opportunities for further optimization of probe design or labeling conditions.

To further confirm target selectivity in a native cellular context, we performed pull-down experiments in PANC1 cells. These experiments successfully enriched Akt2 from complex cellular lysates, validating **probe 3** as a functional chemical tool for target engagement studies. Overall, the reported strategy in this study could be transferred to other Akt isoform-selective molecules to investigate Akt1 or Akt3.

In summary, we were able to design Akt2-selective molecular probes that can be modified with click-chemistry and easily used as a chemical toolbox to analyze Akt2 within complex systems. Their selectivity will allow using those alkyne probes as a powerful handle to analyze and dissect Akt2 in respect of various biological questions. It will help to understand the biology of this specific Akt isoform in health and disease, which ultimately enables the identification of diagnostic-relevant biomarkers and the development of novel therapeutic strategies.

## Methods

### General Chemistry

All reagents and solvents were purchased from Acros, Activate Scientific, Alfa Aesar, Apollo Scientific, Merck, Sigma-Aldrich, TCI Chemicals or VWR and used without further purification. Dry solvents were purchased as anhydrous reagents from commercial suppliers. ^1^H and ^13^C NMR spectra were recorded on a Bruker Avance AV500 (500 MHz and 125 MHz), AV600 (600 MHz and 151 MHz) and AV700 (700 MHz and 176 MHz). ^1^H chemical shifts are reported in *λ* (ppm) as s (singlet), d (doublet), dd (doublet of doublet), t (triplet), q (quartet), m (multiplet) and are referenced to the residual solvent signal: CDCl_3_-*d* (7.26), DMSO-*d_6_* (2.50) or MeOD-*d_4_* (3.34). ^13^C spectra are referenced to residual solvent signal: CDCl_3_-*d* (77.1), DMSO-*d_6_* (39.52) or MeOD-*d_4_* (49.86). High-resolution electrospray ionization mass spectra (ESI-FTMS) were recorded on a Thermo LTQ Orbitrap (high-resolution mass spectrometer from Thermo Electron) coupled to an Accela HPLC system supplied with a Hypersil GOLD column (Thermo Electron). LC-MS (ESI-MS) analysis was performed using Agilent HPLC system (1100 series) or (1260 series) with CC 125/4 Nucleodur C18 gravity column (3 µm) from Macherey Nagel coupled to a Thermo Scientific Finnigan LCQ Advantage Max Ion Trap or an Agilent MSD-iQ mass spectrometer. TLC was carried out on Merck 60 F254 aluminium-backed silica gel plates. Compounds were purified by column chromatography using VWR silica gel (40 - 63 µm particle size) or flash chromatography on a Biotage Isolera One and Büchi Reveleris system using Büchi Reveleris Silica Cartridges (4 - 120 g) monitored by UV at *λ* = 210 nm and 280 nm. Preparative HPLC was conducted on a Büchi Reveleris prep system with a VP 125/21 Nucleodur C18 column from Macherey-Nagel and monitored by UV at *λ* = 210 nm and 254 nm. All final compounds were purified to > 95 % purity as determined by high-performance liquid chromatography (HPLC). Purity was measured using Büchi Reveleris Prep system with UV detection at *λ* = 210 nm (system: Nucleodur C18 gravity column from Macherey Nagel 4 mm x 125 mm, 3 µM, 10 – 100 % MeCN in H_2_O, with 0.2 % TFA, for 15 min at 1.0 mL/min).

*Synthesis of tert-butyl (2-oxo-3-(piperidin-4-yl)-2,3-dihydro-1H-benzo[d]imidazol-5-yl)carbamate* (**S3b**) has been reported.^15^

*Synthesis of tert-butyl 4-(6-acrylamido-2-oxo-2,3-dihydro-1H-benzo[d]imidazol-1-yl)piperidine-1-carboxylate* (**S4**) has been reported.^14^

*Synthesis of tert-butyl (4-((4-(6-((tert-butoxycarbonyl)amino)-2-oxo-2,3-dihydro-1H-benzo[d]imidazol-1-yl)pi-peridin-1-yl)methyl)phenyl)boronic acid* (**1**).

The amine (**S3b**, 1.0 g, 3.5 mmol, 1.0 eq.) and 4-formylphenyl boronic acid (625 mg, 4.2 mmol, 1.2 eq.) were dissolved in dry MeOH (5 mL/mmol). A few drops of Et_3_N and AcOH were added, and the mixture stirred for 4 h at 75 °. Followed by addition of 4.0 eq. NaCNBH_3_ (877 mg, 13.9 mmol) and 12 h stirring at 75 °C. The precipitate was filtered off and the solvent removed under reduced pressure. The crude product was suspended in DCM and washed with aq. NaHCO_3_. A reverse C18 cloumn chromatography (water/MeCN + 0.01 % TFA) yielded the desired product **1** (980 mg, 2.3 mmol, 66 %). **^1^H NMR** (600 MHz, MeOD-*d*_4_) *δ* ppm 1.53 (s, 12 H), 2.09 (d, *J* = 13.39 Hz, 2 H), 2.71 - 2.87 (m, 2 H), 3.27 (t, *J* = 13.02 Hz, 2 H), 3.66 (d, *J* = 11.00 Hz, 2 H), 4.39 (s, 2 H), 4.49 (s, 1 H), 6.83 (d, *J* = 6.60 Hz, 1 H), 6.92 - 6.99 (m, 1 H), 7.55 (d, *J* = 5.69 Hz, 2 H), 7.76 (d, *J* = 7.15 Hz, 1 H); **^13^C NMR** (151 MHz, MeOD-*d*_4_) *δ* ppm 26.07, 27.34, 51.82, 60.35, 79.52, 100.26, 109.22, 112.95, 123.86, 129.13, 129.48, 133.51, 134.03, 154.39, 155.06; **HPLC-MS (ESI):** *m/z* für C_24_H_32_BN_4_O_5_ [M+H]^+^, 466.19 calcd. 466.2 found.

*Synthesis of N-(2-oxo-3-(1-(4-(4,4,5,5-tetramethyl-1,3,2-dioxaborolan-2-yl)benzyl)piperidin-4-yl)-2,3-dihydro-1H-benzo[d]imidazol-5-yl)acrylamide* (**2**).

The secondary amine **S4** (445.8 mg, 1.7 mmol, 1.5 eq) was solved in 2 mL AcOH and 4.5 mL dry DCM. Afterwards 4-(4,4,5,5-tetramethyl-1,3,2-dioxaborolan-2-yl)benzaldehyde (270.2 mg, 1.2 mmol, 1 eq.) was charged with 5 mL dry DCM and added to the solution, followed by stirring at rt for 30 min before it was cooled to 0 °C. The base Et_3_N (2.5 mL) was added slowly over 30 min. After 2 h stirring at rt the reducing agent NaBH(OAC)_3_ was added and everything stirred for additional 19 h. The solution was washed with aq. HCl und extracted with DCM. The collected organic phases were dried over Na_2_SO_4_ and the solvent evaporated. After column chromatography (silica, 1-6.5 % MeOH/DCM + 1 % NH_3_), the desired boronic ester **2** was isolated (180.6 mg, 0.36 mmol, 30 %). **^1^H-NMR** (600 MHz, DMSO-*d*_6_) δ ppm 10.79 (s, 1H), 10.08 (s, 1H), 7.71 (d, *J* = 1.9 Hz, 1H), 7.66 (d, *J* = 7.5 Hz, 2H), 7.39 – 7.35 (m, 2H), 7.26 (dd, *J* = 8.4, 1.8 Hz, 1H), 6.91 (d, *J* = 8.4 Hz, 1H), 6.42 (dd, *J* = 16.9, 10.1 Hz, 1H), 6.27 (dd, *J* = 2.0 Hz, 1H), 5.74 (dd, *J* = 10.1, 2.0 Hz, 1H), 4.14 – 4.05 (m, 1H), 3.56 (s, 2H), 2.95 (d, *J* = 11.1 Hz, 2H), 2.30 (m, 2H), 2.13 – 2.06 (m, 2H), 1.64 (d, *J* = 12.1 Hz, 2H), 1.29 (s, 12H). **^13^C NMR** (151 MHz, DMSO-*d*_6_) δ ppm 162.77, 154.00, 141.90, 134.42, 132.64, 132.03, 129.01, 128.40, 126.35, 124.48, 112.12, 108.64, 100.88, 83.56, 61.95, 52.66, 50.18, 40.92, 24.96, 24.68. **HPLC-MS (ESI):** *m/z* für C_28_H_36_BN_4_O_4_ ([M+H]^+^), 503.28 calcd., 503.4 found.

*Synthesis of tert-butyl (3-(1-(4-(5-((1H-pyrazol-4-yl)methyl)-3-phenylpyridin-2-yl)benzyl)piperidin-4-yl)-2-oxo-2,3-dihydro-1H-benzo[d]imidazol-5-yl)carbamate* (**3**).

Molecule **S22** (100 mg, 0.37 mmol, 1 eq.) and **1** (190 mg, 0.40 mmol, 1.1 eq.) were dissolved in 4 mL 1,4-dioxane/water (5:1) and charged with argon for 5 min. Then K_2_CO_3_ (102 mg, 0.74 mmol, 2 eq.) and the catalyst Pd(PPh_3_)_4_Cl_2_ (13.3 mg, 0.02 mmol, 0.05 eq.) were added. The mixture was heated for 2 h at 130 °C in a microwave reactor. Without extraction the solvent was removed and the crude solid purified with column chromatography (silica, DCM, MeOH + 1 % NH_3_, 1-7.5 %). Product (**3**) was isolated as slightly yellow solid (120 mg, 0.18 mmol, 48 %). **^1^H NMR** (600 MHz, DMSO-*d*_6_) *δ* ppm 1.46 (s, 12 H) 1.60 (d, *J* = 12.10 Hz, 2 H), 2.08 (t, *J* = 11.19 Hz, 2 H), 2.28 (qd, *J* = 12.29, 3.48 Hz, 2 H), 2.91 (d, *J* = 11.19 Hz, 2 H), 3.51 (s, 1 H), 4.04 - 4.13 (m, 1 H), 5.48 (s, 1 H), 6.30 (t, *J* = 2.11 Hz, 1 H), 6.83 (d, *J* = 8.44 Hz, 1 H), 7.12 - 7.15 (m, 2 H), 7.19 - 7.25 (m, 2 H), 7.27 - 7.33 (m, 2 H), 7.50 (dd, *J* = 1.83, 0.55 Hz, 2 H), 7.65 (d, *J* = 2.20 Hz, 2 H), 7.94 - 7.96 (m, 1 H), 8.56 (d, *J* = 2.20 Hz, 2 H), 9.12 - 9.21 (m, 1 H), 10.68 (s, 1 H); **^13^C NMR** (151 MHz, DMSO-*d*_6_) *δ* ppm 28.20, 28.49, 30.77, 35.78, 49.93, 51.70, 52.54, 61.28, 78.69, 105.70, 108.53, 123.42, 127.39, 128.07, 128.41, 129.01, 129.21, 129.51, 130.44, 131.87, 133.09, 135.08, 137.85, 138.13, 138.29, 139.31, 139.42, 147.48, 152.95, 153.98, 155.50, 162.31; **HPLC-MS (ESI):** *m/z* für C_39_H_42_N_7_O_3_ ([M+H^+^]), 656.80 calcd., 656.8 found.

*Synthesis of 1-(1-(4-(5-((1H-pyrazol-4-yl)methyl)-3-phenylpyridin-2-yl)benzyl)piperidin-4-yl)-6-amino-1,3-dihy-dro-2H-benzo[d]imidazol-2-one* (**4**).

Molecule **3** (120 mg, 0.18 mmol, 1 eq.) was dissolved in 6 mL DCM and 2 mL TFA. The mixture was stirred for 16 h at room temperature. The solvent was evaporated und reduced pressure and the crude product was used for further steps without purification. **HRMS (ESI):** *m/z* für C_34_H_34_N_7_O ([M+H^+^]), 556.6940 calcd., 556.6942 found.

*Synthesis of 5-((1H-pyrazol-4-yl)methyl)-2-chloro-3-phenylpyridine* (**S22**) has been reported.^14^

*Synthesis of 6-amino-1-(1-(4-(5-isobutyl-6-oxo-3-phenyl-1,6-dihydropyrazin-2-yl)benzyl)piperidin-4-yl)-1,3-di-hydro-2H-benzo[d]imidazol-2-one* (**14b**) has been reported.^15^

*Synthesis of (Z)-4-((3-(1-(4-(5-isobutyl-6-oxo-3-phenyl-1,6-dihydropyrazin-2-yl)benzyl)piperidin-4-yl)-2-oxo-2,3-dihydro-1H-benzo[d]imidazol-5-yl)amino)-4-oxobut-2-enoic acid* (**5a**).

The amine **14b** (50.0 mg, 0.09 mmol, 1.0 eq.) was dissolve in a mixture of DCM und THF (1:1, 5 mL, 10 mL/mmol) and malonic anhydride (10.7 mg, 0.11 mmol, 1.2 eq.). The solution was stirred for 4 h at rt. After completion the yellow precipitate was filtered off and washed with EtOAc. The desired product was isolated without further purifications steps (**5a**, 52 mg, 0.08 mmol, 89 %). **^1^H-NMR** (600 MHz, DMSO-*d*_6_) *δ* ppm 0.97 (d, *J* = 6.60 Hz, 6 H), 1.99 (d, *J* = 12.10 Hz, 2 H), 2.17 - 2.26 (m, 1 H), 2.60 - 2.69 (m, 4 H), 3.17 - 3.28 (m, 2 H), 3.42 (d, *J* = 11.00 Hz, 2 H), 4.34 (s, 3 H), 6.24 (s, 1 H), 6.35 (d, *J* = 12.10 Hz, 1 H), 6.45 (d, *J* = 12.10 Hz, 1 H), 6.90 - 7.00 (m, 2 H), 7.22 (s, 5 H), 7.41 - 7.45 (m, 2 H), 7.46 - 7.50 (m, 2 H), 7.79 (s, 1 H), 9.75 - 9.85 (m, 1 H), 10.44 (s, 1 H), 10.94 (s, 1 H); **^13^C-NMR** (151 MHz, DMSO-*d*_6_) *δ* ppm 23.06, 26.30, 26.79, 41.37, 48.26, 51.60, 59.33, 101.58, 109.45, 113.97, 116.38, 118.35, 125.48, 127.62, 128.35, 129.66, 129.76, 130.64, 131.28, 131.66, 131.81, 132.34, 154.43, 163.25, 165.50, 167.22, 167.35; **HRMS (ESI):** [R_t_]: 6.67 min, *m/z* für C_37_H_39_N_6_O_5_ ([M+H^+^]), 647.2976 calcd., 647.2975 found.

*Synthesis of (Z)-4-((3-(1-(4-(5-((1H-pyrazol-4-yl)methyl)-3-phenylpyridin-2-yl)benzyl)piperidin-4-yl)-2-oxo-2,3-dihydro-1H-benzo[d]imidazol-5-yl)amino)-4-oxobut-2-enoic acid* (**5b**).

The amine **4** (50.0 mg, 0.09 mmol, 1.0 eq.) was dissolve in a mixture of DCM und THF (1:1, 5 mL, 10 mL/mmol) and malonic anhydride (10.7 mg, 0.11 mmol, 1.2 eq.). The solution was stirred for 6 hours at rt. After completion the yellow precipitate was filtered off and washed with EtOAc. The desired product was isolated without further purifications steps (**5b**, 54 mg, 0.08 mmol, 91 %). **^1^H NMR** (600 MHz, DMSO-*d*_6_) *δ* ppm 1.95 (d, *J* = 12.65 Hz, 2 H), 2.57 - 2.69 (m, 2 H), 3.06 - 3.17 (m, 2 H), 3.21 (s, 1 H), 3.38 - 3.51 (m, 2 H), 3.65 - 3.70 (m, 2 H), 4.25 - 4.34 (m, 2 H), 4.40 (s, 1 H), 5.51 (s, 2 H), 6.27 (s, 1 H), 6.29 - 6.32 (m, 1 H), 6.33 - 6.38 (m, 1 H), 6.41 - 6.48 (m, 1 H), 7.03 - 7.08 (m, 1 H), 7.11 - 7.17 (m, 2 H), 7.29 - 7.34 (m, 3 H), 7.35 - 7.44 (m, 4 H), 7.50 - 7.52 (m, 2 H), 7.94 - 8.00 (m, 1 H), 8.55 - 8.62 (m, 1 H), 10.43 (s, 1 H), 10.92 (s, 1 H), 11.18 (s, 1 H); **^13^C NMR** (151 MHz DMSO-*d*_6_) *δ* ppm 8.60, 25.65, 25.83, 45.71, 47.69, 51.00, 51.80, 55.05, 58.85, 105.71, 115.10, 117.05, 127.57, 128.49, 128.60, 129.28, 130.06, 130.21, 130.49, 130.84, 131.52, 132.43, 135.36, 138.03, 138.92, 139.49, 140.95, 147.57, 153.80, 154.76, 162.75, 162.98 - 163.05, 165.72, 166.43, 166.70; **HPLC-MS (ESI):** *m/z* für C_38_H_36_N_7_O_4_ [M+H]^+^, 654.74 calcd., 654.8 found.

*Synthesis of N^1^-(3-(1-(4-(5-isobutyl-6-oxo-3-phenyl-1,6-dihydropyrazin-2-yl)benzyl)piperidin-4-yl)-2-oxo-2,3-di-hydro-1H-benzo[d]imidazol-5-yl)-N^4^-(prop-2-yn-1-yl)maleamide* (**Probe 1**).

The acid **5b** (**5a**, 20.0 mg, 0.03 mmol, 1.0 eq.) and propargylamine (1.7 mg, 0.03 mmol, 1.0 eq.) were suspended in 2 mL dry THF and cooled to 0 °C. Afterwards 0.05 mL of DCC in DCM (0.05 mmol, 1.5 eq.) were slowly added. The mixture was allowed to stir at rt for 6 h. After completion the solution was washed with NaHCO_3_ solution and extracted with DCM, before the organic phase was dried over NaSO_4_. Column chromatography (silica, DCM, MeOH + 1 % NH_3_) gave the desired product as solid (12 mg, 0.017 mmol 58 %). **^1^H-NMR** (600 MHz, DMSO-*d*_6_) *δ* ppm 0.98 (m, 6 H), 1.96 (d, *J* = 12.10 Hz, 2 H), 2.42 (s, 2 H), 2.57 - 2.70 (m, 2 H), 3.11 - 3.25 (m, 2 H), 3.44 (d, *J* = 10.64 Hz, 2 H), 3.95 (dd, *J* = 5.32, 2.38 Hz, 1 H), 4.27 - 4.38 (m, 3 H), 6.20 - 6.28 (m, 1 H), 6.31 - 6.38 (m, 1 H), 6.89 - 7.01 (m, 2 H), 7.15 - 7.20 (m, 3 H), 7.26 - 7.33 (m, 3 H), 7.36 - 7.44 (m, 4 H), 7.70 (s, 1 H), 7.75 - 7.82 (m, 2 H), 8.57 (s, 1 H), 9.10 (t, *J* = 5.32 Hz, 1 H), 9.75 (s, 1 H), 10.91 (s, 1 H); **^13^C-NMR** (151 MHz, DMSO-*d*_6_) *δ* ppm 8.62, 22.59, 25.48, 26.25, 28.04, 40.90, 45.71, 52.37, 73.42, 80.52, 100.91, 108.68, 112.42, 124.58, 126.91, 127.76, 128.99, 129.17, 129.67, 130.17, 132.37, 132.90, 137.94, 153.98, 157.61, 162.92, 164.49; **HRMS (ESI):** [R_t_]: 6.69 min, *m/z* für C_40_H_42_N_7_O_4_ ([M+H^+^]), 684.3292 calcd., 684.3288 found.

*Synthesis of N^1^-(3-(1-(4-(5-((1H-pyrazol-4-yl)methyl)-3-phenylpyridin-2-yl)benzyl)piperidin-4-yl)-2-oxo-2,3-di-hydro-1H-benzo[d]imidazol-5-yl)-N^4^-(prop-2-yn-1-yl)maleamide* (**Probe 2**).

Molecule **5b** (50 mg, 0.07 mmol, 1.0 eq.) and propargylamine (4.6 mg, 0.08 mmol, 1.1 eq.) were suspended in 5 mL dry THF and cooled to 0 °C. Afterwards 0.15 mL of DCC in DCM (0.15 mmol, 2.0 eq.) were slowly added. The mixture was allowed to stir at rt for 6 h. After completion the solution was washed with NaHCO_3_ solution and extracted with DCM, before the organic phase was dried over NaSO_4_. Column chromatography (silica, DCM, MeOH + 1 % NH_3_) gave the desired product as solid (25 mg, 0.03 mmol 51 %). **^1^H-NMR** (600 MHz, MeOH-*d*_4_) *δ* ppm 1.71 - 1.78 (m, 2 H), 2.17 - 2.25 (m, 2 H), 2.45 - 2.56 (m, 2 H), 2.61 (t, *J* = 2.57 Hz, 1 H), 3.00 - 3.07 (m, 2 H), 3.59 (s, 2 H), 4.05 (d, *J* = 2.57 Hz, 2 H), 4.21 - 4.28 (m, 1 H), 5.52 (s, 2 H), 6.30 (s, 1 H), 6.37 (d, *J* = 14.30 Hz, 2 H), 6.99 - 7.03 (m, 1 H), 7.11 - 7.15 (m, 2 H), 7.22 - 7.31 (m, 9 H), 7.56 - 7.58 (m, 1 H), 7.69 - 7.70 (m, 1 H), 7.71 - 7.74 (m, 1 H), 7.83 - 7.86 (m, 1 H), 8.46 - 8.49 (m, 1 H), 9.39 (s, 1 H), 11.18 (s, 1 H); **^13^C-NMR** (151 MHz, MeOH-*d*_4_) *δ* ppm 29.79, 29.84, 49.72, 52.51, 53.44, 54.16, 63.31, 72.82, 80.20, 104.05, 107.46, 110.53, 115.67, 126.89, 128.82, 129.59, 130.39, 130.73, 130.80, 131.12, 132.09, 132.24, 133.58, 133.72, 134.38, 138.21, 139.06, 139.71, 140.03, 140.60, 141.29, 148.01, 156.75, 158.01, 165.21, 167.25; **HRMS (ESI):** *m/z* für C_41_H_39_N_8_O_3_ ([M+H^+^]), 691.3067 calcd., 691.3063 found.

*Synthesis of 3-((6-chloro-5-phenylpyridin-3-yl)methyl)aniline* (**S20**) has been reported.^14^

*Synthesis of tert-butyl(3-((3-((6-chloro-5-phenylpyridine-3-yl)-methyl)phenyl)amino)-3-oxopropyl)carbamate* (**6**).

The aniline **S20** (371.1 mg, 1.26 mmol, 1 eq.) and *β*-Boc-Ala-OH (238.0 mg, 1.89 mmol, 1.0 eq.) were suspended in 3 mL dry THF and cooled to 0 °C. Afterwards 1.9 mL of DCC in DCM (1.89 mmol, 1.5 eq.) were slowly added. The mixture was allowed to stir at rt for 3 d. After completion the solution was washed with NaHCO_3_ solution and extracted with DCM, before the organic phase was dried over NaSO_4_. Column chromatography (silica, DCM, MeOH + 1 % NH_3_) gave the product **6** as white solid (425 mg, 0.91 mmol 73 %). **^1^H-NMR** (600 MHz, DMSO-*d*_6_) δ ppm 9.88 (s, 1H), 8.33 (d, *J* = 2.4 Hz, 1H), 7.70 (d, *J* = 2.4 Hz, 1H), 7.51 – 7.41 (m, 7H), 7.22 (t, *J* = 7.8 Hz, 1H), 6.98 (d, *J* = 7.6 Hz, 1H), 6.83 (t, *J* = 5.7 Hz 1H), 4.00 (s, 2H), 3.18 (q, 2H), 2.43 (t, *J* = 7.1 Hz, 2H), 1.35 (s, 9H). **^13^C NMR** (151 MHz, DMSO-*d*_6_) δ ppm 169.40, 155.52, 148.52, 146.16, 140.44, 139.50, 136.97, 136.68, 135.77, 129.20, 128.97, 128.41, 128,33 123.51, 119.22, 117.23, 77.59, 36.99, 36.73, 36.46, 28.22; **HPLC-MS (ESI):** *m/z* für C_26_H_29_ClN_3_O_3_ [M+H]^+^, 466.19 calcd. 466.2 found.

*Synthesis of tert-butyl (3-((3-((6-(4-((4-(6-acrylamido-2-oxo-2,3-dihydro-1H-benzo[d]imidazol-1-yl)piperidine-1-yl)methyl)phenyl)-5-phenylpyridine-3-yl)methyl)phenyl)amino)-3-oxopropyl)carbamate* (**7**).

Molecule **6** (210.1 mg, 0.45 mmol, 1 eq.) and **2** (294.5 mg, 0.59 mmol, 1.3 eq.) were dissolved in 4 mL 1,4-dioxane/water (5:1) and charged with argon for 5 min. Then K_2_CO_3_ (124.63 mg, 0.9 mmol, 2 eq.) and the catalyst Pd(PPh_3_)_4_Cl_2_ (52.1 mg, 0.05 mmol, 0.1 eq.) were added. The mixture was heated for 2 h at 130 °C in a microwave reactor. Without extraction the solvent was removed and the crude solid purified with column chromatography (silica, DCM, MeOH + 1 % NH_3_, 1-7.5 %). Product (**7**) was isolated as slightly yellow solid (104.1 mg, 0.26 mmol, 29 %). **^1^H-NMR** (600 MHz, DMSO-*d*_6_) δ ppm 10.79 (s, 1H), 10.07 (s, 1H), 9.89 (s, 1H), 8.56 (d, *J* = 2.1 Hz, 1H), 7.67 (d, *J* = 1.9 Hz, 1H), 7.62 (d, *J* = 2.1 Hz, 1H), 7.54 – 7.51 (m, 1H), 7.49 – 7.45 (m, 1H), 7.33 – 7.18 (m, 9H), 7.18 – 7.13 (m, 2H), 7.05 – 7.01 (m, 1H), 6.91 (d, *J* = 8.4 Hz, 1H), 6.83 (t, *J* = 5.7 Hz, 1H), 6.40 (dd, *J* = 16.9, 10.1 Hz, 1H), 6.24 (dd, *J* = 17.0, 2.0 Hz, 1H), 5.73 (dd, *J* = 10.1, 2.0 Hz, 1H), 4.15 – 4.06 (m, 1H), 4.03 (s, 2H), 3.50 (s, 2H), 3.22 – 3.16 (m, 2H), 2.93 (d, *J* = 11.0 Hz, 2H), 2.43 (t, *J* = 7.1 Hz, 2H), 2.33 – 2.24 (m, 2H), 2.07 (t, *J* = 11.8 Hz, 2H), 1.66 – 1.62 (m, 2H), 1.35 (s, 9H); **^13^C NMR** (151 MHz, DMSO-*d*_6_) δ ppm 169.41, 162.75, 155.52, 154.04, 153.99, 148.33, 140.82, 139.57, 139.48, 138.56, 138.51, 137.67, 135.34, 135.06, 132.64, 132.04, 129.50, 129.29, 128.97, 128.95, 128.36, 128.21, 127.27, 126.33, 124.48, 123.59, 119.29, 117.15, 112.10, 108.62, 100.90, 77.59, 61.53, 52.63, 50.13,40,06 37.68, 36.73, 36.47, 28.22; **HPLC-MS (ESI):** *m/z* für C_48_H_52_N_7_O_5_ [M+H]^+^, 806.4 calcd., 806.4 found.

*Synthesis of N-(3-(1-(4-(5-(3-(3-aminopropanamido)benzyl)-3-phenylpyridine-2-yl)benzyl)piperidine-4-yl)-2-oxo-2,3-dihydro-1H-benzo[d]imidazol-5-yl)acrylamide* (**8**).

The protected intermediate **7** (183.6 mg, 0.23 mmol, 1 eq.) was dissolved in 3 mL DCM and 1 mL TFA. The mixture was stirred for 16 h at room temperature. The solvent was evaporated und reduced pressure. A column chromatography (silica, DCM, MeOH + 1 % NH_3_) yielded the desired product **8** (157.6 mg, 0.22 mmol, 98 %). **^1^H-NMR** (600 MHz, DMSO-*d*_6_) δ ppm 10.89 (s, 1H), 10.16 (s, 1H), 10.11 (s, 1H), 8.59 (d, *J* = 2.1 Hz, 1H), 7.81 (s, 4H), 7.66 (s, 1H), 7.56 (s, 1H), 7.46 (dd, *J* = 8.6, 2.0 Hz, 1H), 7.41 (d, *J* = 12.1 Hz, 2H), 7.35 (d, *J* = 8.1 Hz, 2H), 7.32 – 7.23 (m, 5H), 7.19 – 7.13 (m, 3H), 7.10 – 7.05 (m, 2H), 6.99 (s, 1H), 6.94 (d, *J* = 8.3 Hz, 1H), 6.43 (dd, *J* = 16.9, 10.1 Hz, 1H), 6.23 (dd, *J* = 17.0, 2.0 Hz, 1H), 5.77 – 5.71 (m, 1H), 4.33 (d, *J* = 36.9 Hz, 3H), 4.05 (s, 2H), 3.20 (s, 2H), 3.07 (t, *J* = 6.6 Hz, 2H), 2.71 – 2.57 (m, 4H), 2.00 – 1.87 (m, 2H). **^13^C-NMR** (151 MHz, DMSO-*d*_6_) δ ppm 168.34, 162.84, 153.95, 148.42, 140.87, 139.16, 138.58, 135.86, 135.30, 132.43, 131.94, 130.79, 130.01, 129.34, 129.11, 129.00, 128.45, 127.45, 126.38, 124.64, 123.93, 120.31, 119.40, 118.32, 117.27, 116.33, 114.33, 113.30, 108.84, 61.18, 58.88, 56.02, 51.05, 48.60, 47.54, 40.06, 37.69, 34.92, 33.19, 25.89, 18.56, 15.13. **HPLC-MS (ESI):** *m/z* für C_43_H_44_N_7_O_3_ [M+H]^+^, 706.35 calcd., 706.4 found.

*Synthesis of N-(2-oxo-3-(1-(4-(3-phenyl-5-(3-(3-(prop-2-yn-1-ylamino)propanamido)benzyl)pyridine-2-yl)benzyl)piperidine-4-yl)-2,3-dihydro-1H-benzo[d]imidazol-5-yl)acrylamide* (**Probe 3**).

The primary amine **8** (70.8 mg, 0.1 mmol, 1 eq.) and propargyl bromide (11.93 mg, 0.1 mmol, 1 eq.) were dissolved in dry DMF. Followed by addition of K_2_CO_3_ (27.7 mg, 0.2 mmol, 2 eq.), the mixture was stirred for 12 h at rt. Afterwards the solvent was removed under vacuo the crude mixture purified with column chromatography (silica, DCM, EtOAc/MeOH (3:1), 1-50 %). The final product (**8**) was isolated as white solid (22.6 mg, 0.03 mmol, 30 %). **^1^H-NMR** (700 MHz DMSO-*d*_6_) δ ppm 10.82 (s, 1 H), 10.15 (s, 1 H), 10.11 (s, 1 H), 8.57 (d, *J* = 2.1 Hz, 1 H), 7.73 (s, 1 H), 7.64 – 7.62 (m, 1 H), 7.54 (s, 1 H), 7.49 – 7.45 (m, 1 H), 7.32 – 7.22 (m, 10 H), 7.17 – 7.13 (m, 2 H), 7.06 (d, *J* = 7.8 Hz, 1 H), 6.92 (d, *J* = 8.3 Hz, 1 H), 6.43 (dd, *J* = 16.9, 10.2 Hz, 1 H), 6.23 (dd, *J* = 17.0, 2.0 Hz, 1 H), 5.73 (dd, *J* = 10.1, 2.0 Hz, 1 H), 4.20 – 4.10 (m, 1 H), 4.04 (s, 2 H), 3.81 (s, 2 H), 3.74 (q, *J* = 7.1 Hz, 2 H), 3.60 – 3.56 (m, 2 H), 3.18 – 3.14 (m, 2 H), 3.03 – 2.92 (m, 2 H), 2.69 (t, *J* = 6.9 Hz, 2 H), 2.39 – 2.29 (m, 2 H), 1.78 – 1.59 (m, 2 H), 1.10 (t, *J* = 7.1 Hz, 1 H). **^13^C-NMR** (151 MHz, DMSO-*d*_6_) δ ppm 168.58, 162.80, 153.98, 148.35, 140.92, 139.49, 139.21, 138.54, 135.44, 135.13, 132.61, 132.04, 129.62, 129.30, 129.07, 128.98, 128.39, 127.34, 126.32, 124.52, 123.87, 120.33, 119.37, 118.34, 117.24, 116.35, 114.35, 108.69, 101.00, 78.04, 61.18, 54.93, 52.35, 42.46, 40.06, 37.67, 36.23, 33.17, 33.15, 15.13; **HRMS (ESI):** *m/z* für C_46_H_46_N_7_O_3_ [M+H]^+^, 744.3584 calcd., 744.3587 found.

**Reagents and Materials**. Supplies for the Akt HTRF assay were acquired from CisBio (Bagnols-sur-Cez’e, France). Active Enzymes were purchased from ProQinase (Akt1 (#1379-0000-2), Akt3(#1578-0000-1)), and Thermo Fisher Scientific (Akt2 (#PV3184)). Small volume (25 μL fill volume) white round-bottom 384-well plates were obtained from Greiner Bio-One GmbH (Solingen, Germany).

**Activity-Based Assay**. The biochemical half maximal inhibitory concentrations (IC_50_) were acquired with the STK HTRF KinEASE assay (Cisbio) according to the manufactor’s instructions. Briefly: 5 µL Kinase solution and 2.5 µL inhibitor solution (8 % DMSO in HTRF buffer) were incubated for 1 h before the reaction was started by addition of 2.5 µL starting solution containing ATP and substrate peptide. ATP concentrations were set at their respective *K*_M_ values (80 µM for Akt1, 30 µM for Akt2, and 110 µM for Akt3). The following substrate concentrations were used: 250 µM for Akt1 as well for Akt2, and 1 µM for Akt3. After reaction completion (Akt1: 60 min, Akt2: 20 min, Akt3: 15 min), 10 µL of stop solution were added. The FRET signal was measured with an EnVision plate reader (PerkinElmer, Waltham, MA, US) (*λ* ex 620 nm/ *λ* em 665 nm). The quotient of both intensities was recorded at 8 different inhibitor concentrations and data fit to a Hill 4-parameter equation with Quattro software suite (Quattro Research GmbH, Martinsried, Germany). Each reaction was performed in duplicates and at least three independent determinations of each IC_50_ were made.

**Protein expression, purification and crystallization.** A gene encoding for AKT1 (2-446, E114/115/116A), AKT2 (2-447) including an N-terminal His_6_-tag and a TEV protease recognition site was synthesized by GeneArt AG and cloned into the pIEx/Bac3 expression vector (Merck Millipore) using *NcoI* and *BamHI* restriction sites. The mutations S205T, D262E, K268R, N269D and deletion E267 were introduced by PCR. Transfection, virus generation, amplification and protein expression were carried out in *Spodoptera frugiperda* (Sf9) cells (Thermo Fisher Scientific) following the BacMagic protocol (Merck Millipore). Infected insect cells were grown in Erlenmeyer flasks for 72 hours at 27 °C with shaking at 120 rpm. The cells were harvested by centrifugation at 3,000 g for 20 minutes and washed once with PBS before being flash frozen in liquid nitrogen. Afterwards, cells were thawed and resuspended in lysis buffer [50 mMol/L Tris, 500 mMol/L NaCl, 1 mMol/L DTT, 10 % glycerol, 0.1 % Triton X-100, pH 8.0, EDTA-free protease inhibitor cocktail (Sigma-Aldrich)]. Cells were lysed using a microfluidizer, followed by centrifugation (40,000 g, 1 hour). The supernatant was loaded on a Ni-NTA Superflow Cartridge (Qiagen). Bound protein was eluted in buffer containing 50 mMol/L Tris, 500 mMol/L NaCl, 500 mMol/L imidazole, 1 mMol/L DTT, 10% glycerol, pH 8.0. For cleavage of the His_6_-tag, TEV protease was added to the pooled elution fractions and dialyzed overnight into buffer containing 25 mmol/L Tris, 50 mmol/L NaCl, 1 mmol/L DTT, 5 % glycerol, pH 8.0 at 4 °C. The cleaved protein was further purified by size-exclusion chromatography on a HiLoad 16/60 Superdex 75 pg Column (GE Healthcare) using buffer containing 50 mmol/L HEPES, 200 mMol/L NaCl, 1 mMol/L DTT, 10 % glycerol, pH 7.3. Afterwards, the protein was transferred into storage buffer (25 mMol/L Tris, 100 mMol/L NaCl, 5 mMol/L DTT, 10 % glycerol, pH 7.5) using a Superdex 75 10/300 GL Column (GE Healthcare), concentrated and stored at 80 °C. For crystallization, purified protein at a concentration of 3 mg/mL was incubated with 3 equivalents of inhibitor on ice for 60 minutes. The samples were centrifuged at 20,000 g for 10 minutes before hanging drops were prepared in 15-well crystallization plates (EasyXtal Tool, Qiagen) by mixing protein–ligand complex with reservoir solution (1:1) containing 1.25 mMol/L sodium acetate pH 6, 3.75 mMol/L sodium citrate pH 6.5, and 12 % PEG MME 2000 at 20 °C. Crystals grew within 10 days, which were used for soaking to obtain diffraction-grade crystals and were cryoprotected using 20 % ethylene glycol before they were flash cooled in liquid nitrogen. X-ray diffraction data were collected at the PXII-X10SA beam line of the Swiss Light Source (Paul Scherrer Institute, Villigen, Switzerland) with wavelengths close to 1 Å. The diffraction data were integrated with XDS^30^ (X-ray Detector Software) program package and scaled using the program XSCALE.^31^ The crystal structure was solved by molecular replacement with PHASER using a co-crystal structure of Akt1 in complex with another CAAI as template.^20^ The manual rebuilding of the molecule of the asymmetric unit was performed using the program COOT^32^ and with the help of Dundee PRODRG^33^ server the inhibitor topology files were generated. For multiple cycles of refinement, phenix.refine^34^ was employed and the final structure was evaluated by Ramachandran plot analysis using the server MolProbity.^35,36^ Final validation of the model was performed with the help of the PDB_REDO server.^37^

**Mass-Spectrometry.** We used Akt2 for MS experiments and incubated 10 µM of the protein with 100 µM of the inhibitor in a buffer for 2 h. The samples were analyzed by mass spectrometry using a Thermo Fisher Scientific Ultimate 3000 HPLC system connected with a Thermo Fisher Scientific Velos Pro (2D ion trap). The sample (2 µL) was injected and separated by an AdvanceBio Desalting-RP Cartridge (Agilent Technologies) starting at 95 % solvent A (0.1 % formic acid in water) and 5 % solvent B (0.1 % formic acid in acetonitrile) for 30 seconds, followed by a linear gradient over 2.5 minutes up to 80 % solvent B. A mass range of 700-2000 m/z was scanned and resulting raw data were analyzed with MagTran (v1.02) or ProMass for Xcalibur (v2.8 rev.2 Novatia).

**Cell Lines.** PANC1 cells were a gift from Dr. Jens Siveke (DKFZ Essen). HCC1954 cells were kindly gifted from ifADo (Dortmund). Cells were maintained in DMEM or RPMI-1640 medium (Gibco) supplemented with 10 % fetale bovine serum (FBS, PAN-Biotech) and 1 % penicillin-streptomycin (Gibco). All cells were cultured at 37 °C at 5 % CO_2_.

**Generation of HEK293T Akt2 KO.** Single-guide RNAs (sgRNAs) targeting human Akt2 were designed using CRISPick, CHOPCHOP, and the Invitrogen TrueDesign Genome Editor to maximize predicted on-target activity and minimize off-target effects. Complementary oligonucleotides were phosphorylated using T4 polynucleotide kinase (37 °C for 30 min; enzyme inactivation at 65 °C for 20 min) and annealed by heating to 95 °C followed by gradual cooling to 4 °C. The annealed sgRNA duplexes was ligated into BbsI-digested pSpCas9(BB)-2A-GFP vector. Recombinant plasmids were transformed into chemically competent *E. coli* XL10-Gold cells, and correct sgRNA insertion was confirmed by Sanger sequencing after plasmid purification.

For genome editing, HEK293T cells were seeded at 5 × 10^5 cells per well in 24-well plates 24 h prior to transfection and transfected with the Akt2-targeting pSpCas9(BB)-2A-GFP construct using Lipofectamine™ 3000 (Thermo Fisher Scientific) according to the manufacturer’s protocol. Twenty-four hours post-transfection, cells were prepared for fluorescence-activated cell sorting (FACS). Cells were detached using trypsin, resuspended in filtered culture medium, and passed through a cell strainer to obtain a single-cell suspension. GFP-positive cells were sorted using a SH800S Cell Sorter (Sony), and single GFP-positive cells were deposited into individual wells of 96-well plates containing culture medium.

Clonal populations were expanded for up to three weeks with regular medium replacement. Genomic DNA was isolated using the QIAamp DNA Mini Kit (Qiagen), and the targeted Akt2 locus was amplified by PCR using gene-specific primers. Genome editing was confirmed by Sanger sequencing. Validated monoclonal Akt2-edited cell lines were cryopreserved at 1 × 10^6 cells/mL using controlled-rate freezing and stored at −150 °C for longterm preservation.

**Cell treatment and lysis.** Cells were seeded into six-well tissue culture plates (Sarstedt) (HCC1954: 500.000 cells/well, PANC1: 250.000 cells/well; HEK293: 250.000 cells/well). After 24-48 hours incubation, cells were treated with various concentrations of probes or DMSO and incubated for additional 20 h before cells were washed twice with ice-cold PBS. Cell lysis was initiated by addition of 100 μL lysis buffer (DPBS, 1 % Triton X-100) per well supplemented with phosphatase and protease inhibitor cocktails (Sigma) followed by incubation on ice for 20 min. Cells were then harvested by scraping. Whole cell lysates were cleared by centrifugation at 14,000 x g/4 °C for 15 min and transferred into fresh, precooled microcentrifuge tubes. Protein concentrations were determined using the Pierce BCA protein assay (Thermo) as per manufacturer’s instructions and normalized to 1 mg/mL.

**Cell lysate labeling.** Soluble proteome samples from untreated cells were obtained as described above and normalized to a concentration of 1 mg/mL in a volume of 100 µL. Lysates were then treated with inhibitor or probe (0.1 µL of 1000x stock in DMSO) mixed by agitation and incubated for the indicated times at 21 °C. For the competition experiment with borussertib 3 h treatment with inhibitor was followed by 1.5 h probe treatment.

**Click Chemistry.** Click Chemistry was then performed with the normalized labelled, or DMSO treated proteome samples in a volume of 50 µL. On each sample a final concentration of either 25 µM Cy5.5- or Biotin-PEG3-azide plus the reagents to catalyse the reaction 1 mM Tris(2-carboxyethyl)phosphine (TCEP, Sigma-Aldrich), 100 µM Tris[(1-benzyl-1H-1,2,3-triazol-4-yl)methyl]amine (TBTA, Sigma-Aldrich) and 1 mM CuSO_4_ were added. In the control experiments the copper was replaced by water. If the samples were treated with fluorophore, they were incubated at room temperature by light agitation for 1 h. In case biotin-azide was used, the samples were incubated at 4 °C over night. After reaction completion the samples treated with copper were centrifuged at 6400 g at 4 °C for 5 min and the supernatant was removed. To each protein pellet 500 µL ice cold methanol was added and thoroughly vortexed. After centrifugation at 6400 g for 5 min the methanol was removed, and this washing step repeated for two more times. For solubilization a maximum of 50 µL of 1 % SDS in PBS was added and the tubes sonicated 1-5 min until the pellet dissolved properly. Biotin samples were then used for further pull-down experiments, whereas for western blot analysis the samples were treated with 4x SDS loading buffer and used as described below.

**Pull-down.** For the protein enrichment magnetic streptavidin beads (Dynabeads™ MyOne™ Streptavidin C1, Thermo) were used. For 50 µL of biotin-labeled sample 30 µL of beads were prepared through washing them with 1 mL of TBS-T. After vortexing them and separating the beads for 1 min on the magnet the solution was removed entirely. This step was repeated three times in total. Then 50 µL of sample was added to the beads and incubated at 4 °C over night with over-head rotation. On the next day the samples were separated over sitting for 1 min on the magnet. The supernatant was stored on ice if needed for western blot analysis and represents the not enriched fraction. Followed by washing the beads with 1 mL PBS three times and thorough vortexing in between. To release the proteins for western blot analysis the beads were boiled in 50 µL 0.1 % SDS in PBS at 95 °C for 5 min. After separation over a magnet the supernatant was transferred to a new vial and 4x SDS loading buffer was added as well as to the not-enriched fraction from before.

**Fluorescence Microscopy.** Cells were seeded onto glass coverslips coated with 0.01 % poly-L-lysine (Sigma-Aldrich, P4832, 1:10 dilution). Coated coverslips were placed in 6-well plates, and cells were plated at a density of 1.5 × 10⁵ cells per well. The following day, cells were treated with the indicated inhibitor and incubated overnight. On the next day, the inhibitor was washed out by washing the wells three times with phosphate-buffered saline (PBS), followed by three medium changes at 10 min intervals. Cells were fixed with 3.7 % formaldehyde for 15 min at room temperature and washed three times with PBS. Permeabilization was performed using PBS containing 0.25 % Triton X-100, followed by blocking with 3 % bovine serum albumin (BSA) prepared in PBS containing 0.25 % Triton X-100. After blocking, the inhibitor in cells was labeled with Cy3-azide using the Click-iT® Cell Reaction Buffer Kit (Thermo Fisher Scientific, Catalog no. C10269) according to the manufacturer’s instructions. Cells were subsequently washed five times with PBS containing 0.25 % Triton X-100 and counter-stained with DAPI (1:1000 dilution) for 1 h at room temperature. Coverslips were mounted onto glass slides using Mowiol 4-88 mounting medium and allowed to dry overnight at room temperature. Fluorescence images were acquired using an Olympus IX81 inverted widefield fluorescence microscope equipped with a Lumencor Spectra X light source and a Hamamatsu ORCA-R² CCD camera, using a 20× UPLSAPO air objective.GFP-Akt2, Cy3, and DAPI fluorescence were detected using the appropriate filter sets, with exposure times of 50 ms, 200 ms, and 10 ms, respectively. Images were acquired using Olympus cellSens Dimension software and processed using Fiji/ImageJ (ImageJ 2.16.0). For image presentation, linear adjustments of brightness and contrast were applied equally across all conditions. Quantitative image analysis was performed using CellProfiler by segmenting nuclei based on DAPI staining and identifying corresponding cellular regions, followed by measurement of mean fluorescence intensity of Cy3-**probe 3** channel. CellProfiler analysis pipelines were executed using identical settings for all experimental conditions.

**Western Blot analysis.** Equal amounts of protein (10-13 µL) were separated by a 12 % SDS-PAGE gel and transferred to Immobilon-FL PVDF membranes (Merck Millipore) using Pierce™ 1-step transfer buffer (Thermo) and the Pierce™ Power Blotter (Thermo). Membranes were washed for 5 min with ddH_2_O, blocked with

Intercept®Blocking Buffer TBS (Li-Cor) for 1 h at room temperature. Biotin labeled samples were incubated with SA-DyLight in Intercept®Blocking Buffer TBS for another hour at rt and then washed three times with TBS-T (50 mM Tris, 150 mM NaCl, 0.05 % Tween 20, pH 7.4) for 5 min. Followed by incubation with primary antibodies diluted in Intercept®Blocking Buffer TBS overnight at 4 °C with gentle agitation. On the next day, the membranes were washed three times with TBS-T for 5 min before being incubated with secondary antibody diluted in Intercept®Blocking Buffer TBS for 1 h at room temperature. Finally, the membranes were washed three times for 5 min with TBS-T and then scanned using an Odyssey®CLx imaging system (Li-Cor).

**Antibodies.** Anti-tAkt1 (CST, cat. No. 2938, 1:1000), anti-tAkt2 (CST, cat. No. 3063, 1:1000), anti-pAkt1 (CST, cat. No. 9018, 1:1000), anti-pAkt2 (CST, cat. No. 8599, 1:1000), anti-rabbit IgG (H+L) (DyLight™ 800 4X PEG Conjugate) (CST, cat. No. 5151, 1:15000), anti-HSP90 (CST, cat. No. 4877, 1:15000), Streptavidin Protein-DyLight™ 650 (Thermo, cat. No. 84547, 1:5000).

## Data Availability

The data supporting the findings of this study are available in the paper and its Supplementary Information. Source Data are provided with this paper. The crystal structure data generated in this study have been deposited in the PDB database under accession code 29mj [http://doi.org/10.2210/pdb29mj/pdb]. Already reported Structures are deposited under following accession codes: 6s9w.

## Acknowledgements

L.Q. is grateful for a scholarship by the German Academic Scholarship Foundation and for a Walter-Benjamin Fellowship by the German Research Council. This work is supported by funds provided by the German Research Foundation (DFG, INST 212/474-1 FUGG), the State of North Rhine-Westphalia (NRW), the European Union (European Regional Development Fund: Investing In Your Future) (EFRE-800400), Drug Discovery Hub Dortmund (DDHD), the German Cancer Aid ((Deutsche Krebshilfe), Targeting Transcriptional Addiction in Cancer (TACTIC, 70115201), as part of the preclinical cancer drug development network (preCDD)), the “Netzwerke 2021” program, an initiative of the Ministry of Culture and Science of the State of North Rhine-Westphalia (CANcer TARgeting, NW21-062C).

## Author Contributions

D.R. and L.Q. are responsible for initiating and supervising the project. L.Q. designed and together with M.B. synthesized the probes. S.B. synthesized the pan-Akt biotin-probe. L.Q. carried out the biochemical experiments and performed cell biology studies. T.K. did pull-down experiments. D.D. did the fluorescence microscopy and HEK293 cell experiments. Leif D. advised on the cell and microscopy experiments. F.S. generated the knock-out cell line. J.N, Laura D., M.P.M. carried out the structural biology studies. The manuscript was written through contributions of all authors. All authors have given approval to the final version of the manuscript.

## Competing Interests

The authors declare no competing interests.

## Notes

http://doi.org/10.2210/pdb29mj/pdb

### Competing Interest Statement

The authors have declared no competing interest.

## References

1 Franke, T. F. PI3K/Akt: getting it right matters. Oncogene 27, 6473–6488 (2008). 10.1038/onc.2008.313

2 Hers, I., Vincent, E. E. & Tavare, J. M. Akt signalling in health and disease. Cellular signalling 23, 1515–1527 (2011). 10.1016/j.cellsig.2011.05.004

3 Smit, D. J. & Jucker, M. AKT Isoforms as a Target in Cancer and Immunotherapy. Curr Top Microbiol Immunol 436, 409–436 (2022). 10.1007/978-3-031-06566-8_18

4 Gonzalez, E. & McGraw, T. E. The Akt kinases: isoform specificity in metabolism and cancer. Cell cycle 8, 2502–2508 (2009). 10.4161/cc.8.16.9335

5 Kupriyanova, T. A. & Kandror, K. V. Akt-2 binds to Glut4-containing vesicles and phosphorylates their component proteins in response to insulin. The Journal of biological chemistry 274, 1458–1464 (1999).

6 Santi, S. A. & Lee, H. The Akt isoforms are present at distinct subcellular locations. American journal of physiology. Cell physiology 298, C580–591 (2010). 10.1152/ajpcell.00375.2009

7 Cho, H. et al. Insulin resistance and a diabetes mellitus-like syndrome in mice lacking the protein kinase Akt2 (PKB beta). Science 292, 1728–1731 (2001). 10.1126/science.292.5522.1728

8 Katome, T. et al. Use of RNA interference-mediated gene silencing and adenoviral overexpression to elucidate the roles of AKT/protein kinase B isoforms in insulin actions. The Journal of biological chemistry 278, 28312–28323 (2003). 10.1074/jbc.M302094200

9 Tschopp, O. et al. Essential role of protein kinase B gamma (PKB gamma/Akt3) in postnatal brain development but not in glucose homeostasis. Development 132, 2943–2954 (2005). 10.1242/dev.01864

10 Dummler, B. et al. Life with a single isoform of Akt: mice lacking Akt2 and Akt3 are viable but display impaired glucose homeostasis and growth deficiencies. Molecular and cellular biology 26, 8042–8051 (2006). 10.1128/MCB.00722-06

11 Linnerth-Petrik, N. M., Santry, L. A., Petrik, J. J. & Wootton, S. K. Opposing functions of Akt isoforms in lung tumor initiation and progression. PloS one 9, e94595 (2014). 10.1371/journal.pone.0094595

12 Riggio, M. et al. AKT1 and AKT2 isoforms play distinct roles during breast cancer progression through the regulation of specific downstream proteins. Scientific reports 7, 44244 (2017). 10.1038/srep44244

13 Stockwell, B. R. Exploring biology with small organic molecules. Nature 432, 846–854 (2004). 10.1038/nature03196

14 Quambusch, L. et al. Cellular model system to dissect the isoform-selectivity of Akt inhibitors. Nature Communications 12, 5297 (2021). 10.1038/s41467-021-25512-8

15 Quambusch, L. et al. Covalent-Allosteric Inhibitors to Achieve Akt Isoform-Selectivity. Angewandte Chemie 58, 18823–18829 (2019). 10.1002/anie.201909857

16 Adam, G. C., Sorensen, E. J. & Cravatt, B. F. Chemical strategies for functional proteomics. Molecular & Cellular Proteomics 1, 781–790 (2002).

17 Speers, A. E. & Cravatt, B. F. Profiling enzyme activities in vivo using click chemistry methods. Chemistry & biology 11, 535–546 (2004).

18 Page, N. et al. Identification and development of a subtype-selective allosteric AKT inhibitor suitable for clinical development. Scientific reports 12, 15715 (2022). 10.1038/s41598-022-20208-

19 Craven, G. B. et al. Mutant-selective AKT inhibition through lysine targeting and neo-zinc chelation. Nature 637, 205–214 (2025). 10.1038/s41586-024-08176-4

20 Weisner, J. et al. Preclinical Efficacy of Covalent-Allosteric AKT Inhibitor Borussertib in Combination with Trametinib in KRAS-Mutant Pancreatic and Colorectal Cancer. Cancer research 79, 2367–2378 (2019). 10.1158/0008-5472.CAN-18-2861

21 Uhlenbrock, N. et al. Structural and chemical insights into the covalent-allosteric inhibition of the protein kinase Akt. Chemical Science 10, 3573–3585 (2019). 10.1039/C8SC05212C

22 Rauh, D., Reyda, S., Klebe, G. & Stubbs, M. T. Trypsin mutants for structure-based drug design: expression, refolding and crystallisation. Biol Chem 383, 1309–1314 (2002). 10.1515/bc.2002.148

23 Lapierre, J.-M. et al. Discovery of 3-(3-(4-(1-Aminocyclobutyl)phenyl)-5-phenyl-3H-imidazo[4,5-b]pyridin-2-yl)pyridin-2-amine (ARQ 092): An Orally Bioavailable, Selective, and Potent Allosteric AKT Inhibitor. Journal of medicinal chemistry 59, 6455–6469 (2016). 10.1021/acs.jmedchem.6b00619

24 Huang, S. et al. Determining Michael acceptor reactivity from kinetic, mechanistic, and computational analysis for the base-catalyzed thiol-Michael reaction. Polymer Chemistry 12, 3619–3628 (2021). 10.1039/D1PY00363A

25 Samavarchi-Tehrani, P., Samson, R. & Gingras, A.-C. Proximity Dependent Biotinylation: Key Enzymes and Adaptation to Proteomics Approaches*. Molecular & Cellular Proteomics 19, 757–773 (2020). 10.1074/mcp.R120.001941

26 Vasudevan, K. M. et al. AKT-Independent Signaling Downstream of Oncogenic PIK3CA Mutations in Human Cancer. Cancer cell 16, 21–32 (2009). 10.1016/j.ccr.2009.04.012

27 Lanning, B. R. et al. A road map to evaluate the proteome-wide selectivity of covalent kinase inhibitors. Nat Chem Biol 10, 760–767 (2014). 10.1038/nchembio.1582

28 Blagg, J. & Workman, P. Choose and Use Your Chemical Probe Wisely to Explore Cancer Biology. Cancer cell 32, 9–25 (2017). 10.1016/j.ccell.2017.06.005

29 Troup, R. I., Fallan, C. & Baud, M. G. J. Current strategies for the design of PROTAC linkers: a critical review. Explor Target Antitumor Ther 1, 273–312 (2020). 10.37349/etat.2020.00018

30 Kabsch, W. XDS. Acta Crystallogr D Biol Crystallogr 66, 125–132 (2010). 10.1107/S0907444909047337

31 Kabsch, W. Automatic Processing of Rotation Diffraction Data from Crystals of Initially Unknown Symmetry and Cell Constants. J Appl Crystallogr 26, 795–800 (1993). Doi 10.1107/S0021889893005588

32 Emsley, P. & Cowtan, K. Coot: model-building tools for molecular graphics. Acta Crystallogr D 60, 2126–2132 (2004). 10.1107/S0907444904019158

33 Schuttelkopf, A. W. & van Aalten, D. M. F. PRODRG: a tool for high-throughput crystallography of protein-ligand complexes. Acta Crystallographica Section D-Structural Biology 60, 1355–1363 (2004). 10.1107/S0907444904011679

34 Afonine, P. V. et al. Towards automated crystallographic structure refinement with phenix. refine. Acta Crystallographica Section D: Biological Crystallography 68, 352–367 (2012).

35 Chen, V. B. et al. MolProbity: all-atom structure validation for macromolecular crystallography. Acta Crystallographica Section D-Structural Biology 66, 12–21 (2010). 10.1107/S0907444909042073

36 Adams, P. D. et al. PHENIX: a comprehensive Python-based system for macromolecular structure solution. Acta Crystallogr D 66, 213–221 (2010). 10.1107/S0907444909052925

37 Joosten, R. P., Long, F., Murshudov, G. N. & Perrakis, A. The PDB_REDO server for macromolecular structure model optimization. Iucrj 1, 213–220 (2014). 10.1107/S2052252514009324

