## Supplementary Information for "Selective covalent-allosteric tools to dissect Akt2"

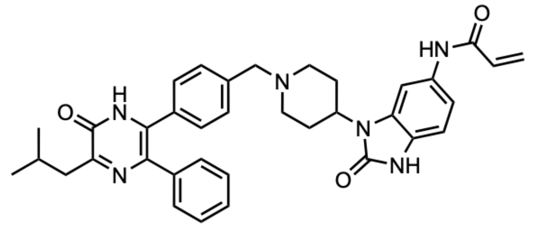

**16b**

$IC_{50}$ (HTRF) /nM
**Akt1:** 1140
**Akt2:** 120
**Akt3:** 16320

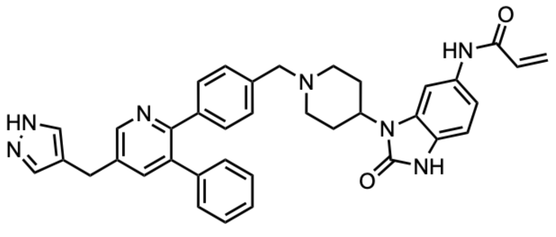

**14**

$IC_{50}$ (HTRF) /nM
**Akt1:** 530
**Akt2:** 25
**Akt3:** 360

**SI-Fig. 1:** Structures and reported $IC_{50}$-values of Akt2-selective CAAIs. [1,2]

**Table S1**: Data collection and refinement statistics for full-length Akt1 mutant in complex with allosteric Akt inhibitor miransertib.

| | Akt1 mutant in complex with miransertib |
|---|---|
| **PDB-ID** | 29MJ |
| **Data collection** | EIGER2 16M @ PXII X10SA SLS |
| Space group | P 21 21 21 |
| Cell dimensions<br>a, b, c [Å]<br>*α, β, γ* [°] | <br>44.80, 164.25, 188.96<br>90.00, 90.00, 90.00 |
| Resolution [Å] | 47.37-2.50<br>(2.6 - 2.5) |
| $R_{meas}$ [%] | 12.2 (154) |
| $I / \sigma I$ | 1.58 (13.99) |
| Completeness [%] | 99.91 (99.94) |
| $CC_{1/2}$ | 0.999 (0.741) |
| Redundancy | 13.0 (13.6) |
| **Refinement** | |
| Resolution [Å] | 47.37-2.50 |
| No. Reflections ($R_{work}$/$R_{free}$) | 49435/2470 |
| $R_{work}$ / $R_{free}$ | 0.228/0.264 |
| No. atoms | 8388 |
| Protein | 8148 |
| Ligands | 115 |
| Water | 125 |
| *B*-factors (Å$^2$) | 75.48 |
| Protein | 75.91 |
| Ligand | 60.34 |
| Water | 61.77 |
| RMS deviations<br>Bond lengths [Å]<br>Bond angles [°] | <br>0.003<br>0.75 |
| Ramachandran [%]<br>Favored<br>Allowed<br>Outliers | <br>96.27<br>3.73<br>0.00 |

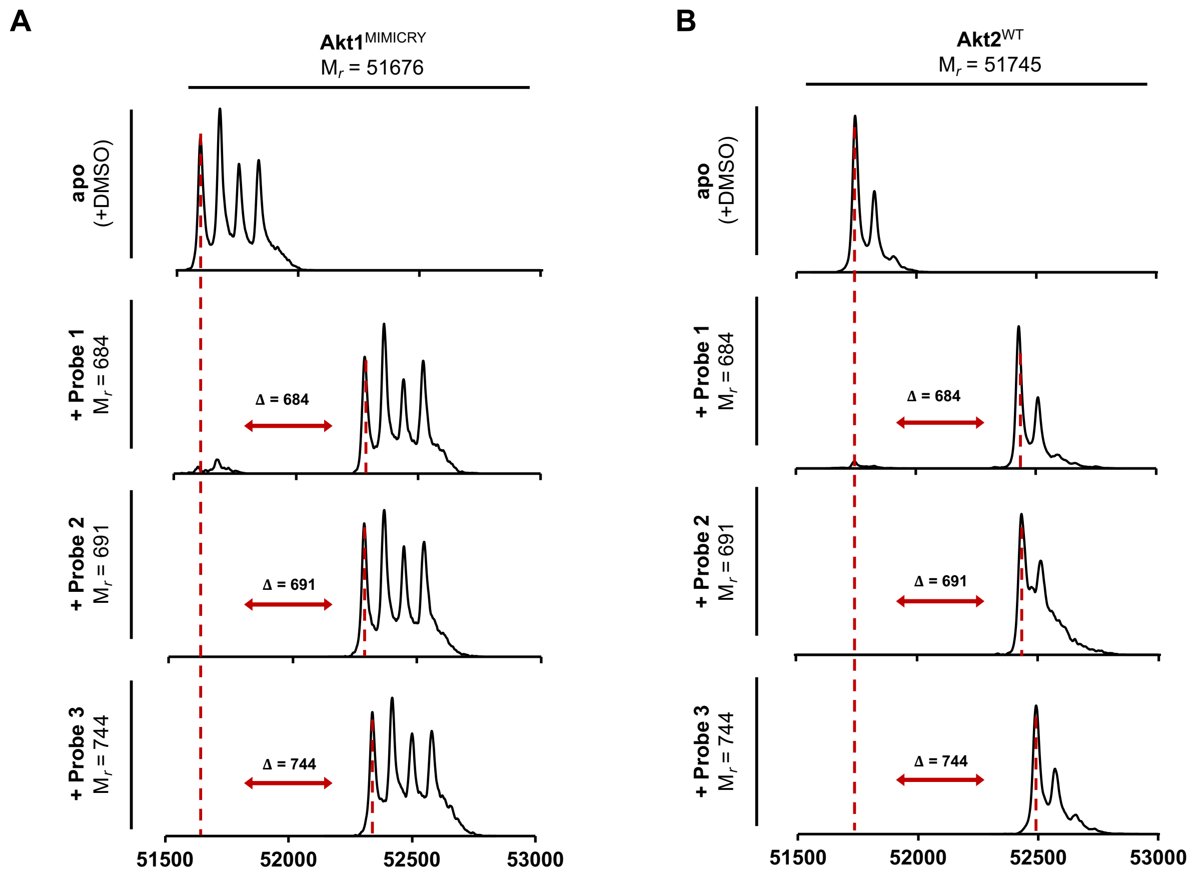


**SI-Fig. 2: Protein mass spectrometry analysis of Probes with the Akt.** Deconvoluted mass spectra of Akt1$^{MIMICRY}$ (A) and Akt2$^{WT}$ (B) after incubation with DMSO (apo) and selected covalent-allosteric Akt probes. All tested molecules show mass differences according to a mono-labeling of the protein. Mass spectra were recorded using denaturing conditions.

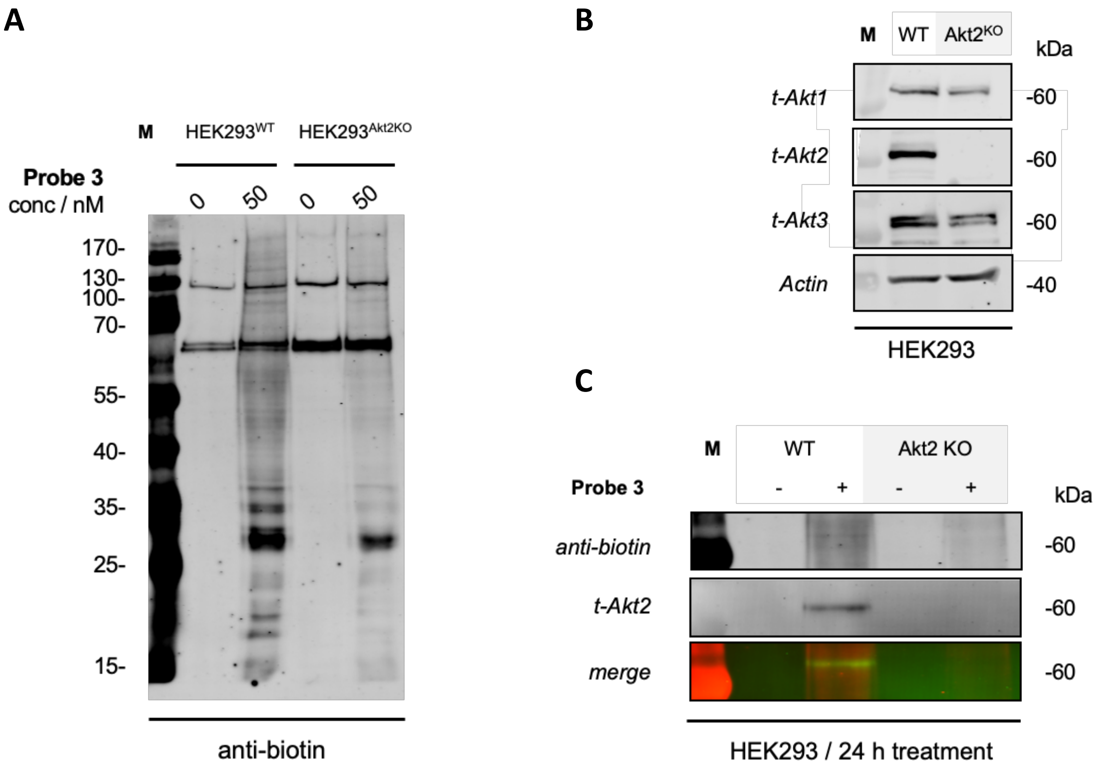


**SI-Fig. 3: Immunoblot of Pulldown experiment in HEK293 wildtype and Akt2 Knock-out cell line.** A: Overview of a biotin-stained immunoblot of HEK293 biotin-streptavidin pull-down experiments treated with or without **probe3** after modification with biotin-PEG3-azide *via* click-chemistry; B: Immunoblot staining for total Akt isoforms level in HEK293 wildtype and Akt2 knock-out cell lines; C: Compiled immunoblot of the experiment shown in A but stained with t-Akt2 antibody to assess the isoform-selective labeling of the pull-down in both HEK293 cell lines.

**References**

1 Quambusch, L. *et al.* Covalent-Allosteric Inhibitors to Achieve Akt Isoform-Selectivity. *Angewandte Chemie* **58**, 18823-18829 (2019). https://doi.org:10.1002/anie.201909857

2 Quambusch, L. *et al.* Cellular model system to dissect the isoform-selectivity of Akt inhibitors. *Nature Communications* **12**, 5297 (2021). https://doi.org:10.1038/s41467-021-25512-8
